# Transient states during the annealing of mismatched and bulged oligonucleotides

**DOI:** 10.1101/2023.09.27.559754

**Authors:** Marco Todisco, Dian Ding, Jack W. Szostak

## Abstract

Oligonucleotide hybridization is crucial in various biological, prebiotic and nanotechnological processes, including gene regulation, non-enzymatic primer extension and DNA nanodevice assembly. Although extensive research has focused on the thermodynamics and kinetics of nucleic acid hybridization, the behavior of complex mixtures and the outcome of competition for target binding remain less well understood. In this study, we investigate the impact of mismatches and bulges in a 12bp DNA or RNA duplex on its association (*k*_*on*_) and dissociation (*k*_*off*_) kinetics. We find that such defects have relatively small effects on the association kinetics, while the dissociation kinetics vary in a position-dependent manner by up to 6 orders of magnitude. Building upon this observation, we explored a competition scenario involving multiple oligonucleotides, and observed a transient low specificity of probe hybridization to fully vs. partially complementary targets in solution. We characterize these long-lived metastable states and their evolution toward equilibrium, and show that sufficiently long-lived mis-paired duplexes can serve as substrates for prebiotically relevant chemical copying reactions. Our results suggest that transient low accuracy states may spontaneously emerge within all complex nucleic acid systems comprising a large enough number of competing strands, with potential repercussions for gene regulation in the realm of modern biology and the prebiotic preservation of genetic information.

## INTRODUCTION

Precise target recognition in nucleic acid hybridization is important for gene regulation (1), PCR (2), *in-situ* hybridization (3, 4), proximity ligation assay (5), non-enzymatic template copying (6), assembly of nanodevices (7, 8) and supramolecular structures (9, 10). Failure in nucleic acid target recognition can lead to misregulation of biological processes (11, 12), amplification of erroneous sequences, and loss of information. Nucleic acid thermodynamics has been extensively studied for sets of paired oligonucleotides, leading to the nearest neighbor (NN) model for the prediction of hybridization energy (13–15). The NN model is a reliable and widely used resource for the design of optimal sequences that minimize mispairings and cross-interactions within sets of oligonucleotides. Over the years the model has been expanded to include the description of a wide range of possible pairing configurations of multiple strands, involving dangling ends, coaxial stacking, loops, bulges, and mismatches. Systematic studies of internal mismatches and bulges have found them to be destabilizing both in a sequence context-dependent and position-dependent manner, with larger effects at internal locations (16–18). One of the major limitations of the NN model is its inherent restriction to the equilibrium description of the system, while neglecting the out-of-equilibrium dynamics, which can be dramatically different (19–22). Specifically, duplexes that are not expected at equilibrium may initially be present at high concentrations when multiple strands are mixed, with abundances that depend on the relative hybridization rates of the various competing species. The available literature on the effect of mismatches on hybridization rates is mostly limited to measurements of surface-immobilized oligonucleotides (23–25), showing a position-dependent and sequence context-dependent effect that usually does not affect the hybridization kinetics (*k*_*on*_) by more than one order of magnitude. In contrast Cisse and colleagues (26) found that hybridization of short fluorescently labeled DNA or RNA oligonucleotides enclosed in 200 nm diameter vesicles slows down by up to two orders of magnitude when mismatches are introduced. The authors proposed an empirical rule to explain their results, suggesting that seven contiguous base pairs are required for the fast hybridization of short DNA or RNA oligonucleotides, leading to the formulation of the so-called “rule of seven” (26).

Comparable hybridization rates are a prerequisite for the creation of metastable states that are significantly different from the final equilibrium state (20). A direct consequence of the dramatic effect of mismatches in hybridization rates as determined by Cisse and colleagues would be that almost no mismatched duplexes are expected in solution at any time if these can be outcompeted by the formation of perfectly paired duplexes. If this is true, the equilibrium description is always well-suited for such systems and cross-interactions are negligible. In this work, we present a systematic study on the effect of single mismatches, double mismatches, and bulges (collectively referred to as “defects”) on the association and dissociation kinetics of short complementary DNA or RNA strands in solution. To evaluate the kinetic effects of these defects, we used a combination of fluorescence-based techniques that exploit the emission of the adenine analogue 2-aminopurine. Finally, we tested the null hypothesis of equivalence between the equilibrium and out-of-equilibrium distribution of defect-containing and perfectly paired duplexes in competing mixtures. Our results reveal the formation of transient low accuracy states during oligonucleotide hybridization in complex mixtures. In such metastable states, defect-containing duplexes that are not present at equilibrium can behave as substrates for subsequent reactions. Understanding the kinetics of nucleic acid annealing in complex mixtures enhances our ability to predict and manipulate nucleic acid hybridization processes.

## MATERIALS AND METHODS

### General

All DNA oligonucleotides were acquired from IDT with standard desalting and were used as received. RNA oligonucleotides were synthesized in-house. Phosphoramidites were acquired from Chemgenes (Wilmington, MA). Controlled pore glass (CPG) columns and all reagents for oligonucleotide synthesis and purification were acquired from Glen Research (Sterling, MA), while reagents used for cleavage and deprotection were acquired from Sigma-Aldrich (St. Louis, MO). All kinetics and competition experiments were performed in 1 M NaCl, 10 mM Tris-HCl buffer pH 8.0 unless otherwise specified. Primer extension and ligation reactions were performed in 100 mM MgCl_2_, 200 mM Tris-HCl buffer pH 8.0. Buffer mixtures were prepared using Tris (1M), sodium chloride (5M) and magnesium chloride (1M) stock solutions from Invitrogen and pH-adjusted using HCl from Sigma-Aldrich. All fluorescence spectroscopy measurements were performed by tracking the fluorescence emission of 2-aminopurine (2Ap), a fluorescent base analogue of adenine whose quantum yield is reduced when in a double helix (27).

### Solid phase synthesis of RNA oligonucleotides

RNA oligonucleotides were synthesized on an Expedite 8909 through conventional solid-phase synthesis, leaving the final 5′-DMT group on. Oligonucleotides synthesized in this way were cleaved from the solid support using AMA (1.2 ml, 1:1 mixture of 40% w/w methylamine aqueous solution and 28% w/w NH_4_OH aqueous solution) for 30 minutes. After cleavage, the oligonucleotides were incubated at 65°C for 20 minutes for deprotection. The deprotected oligonucleotides were dried for 30 minutes at 40°C in a speed-vac and then lyophilized overnight. The RNA powder was resuspended in DMSO (115 µl), TEA (60 µl) and TEA.3HF (75 µl), then incubated at 65°C for 2.5 hours to remove the TBDMS protecting groups. The resulting oligonucleotides were purified and deprotected from the 5′-DMT groups using GlenPak columns (Glen Research). Oligonucleotide concentrations were determined spectrophotometrically using a NanoDrop 2000 (Thermo Scientific) using the extinction coefficients computed through the IDT oligo analyzer (28).

### Synthesis of reactive species

The imidazolium-bridged C*C dimer and the 5′-2-aminoimidazole-activated pCGCA (*CGCA) were synthesized as previously described (29). Briefly, the imidazolium-bridged C*C dimer was prepared by mixing cytidine monophosphate (1 equivalent) and 2-aminoimidazole (2AI) (0.45 equivalents) in dry dimethyl sulfoxide (DMSO). To this mixture, triethylamine (TEA) (20 equivalents), triphenylphosphine (TPP) (10 equivalents), and 2,2-dipyridyldisulfide (DPDS) (10 equivalents) were added. After a 30-minute incubation, the resulting product was precipitated using acetone with sodium perchlorate. The precipitate was then washed twice with a mixture of acetone and diethyl ether (1:1, v/v) and dried under vacuum. Subsequently, the dry pellet was resuspended in 20 mM triethylamine-bicarbonate buffer (TEAB), pH 8.0, and purified using reverse-phase flash chromatography on a 50 g C18Aq column over 12 column volumes (CVs) of 0-10% acetonitrile in 2 mM TEAB buffer (pH 8.0).

For the activated tetranucleotide *CGCA, the 5′-monophosphate oligonucleotide was activated by mixing 2AI (40 equivalents), TPP (40 equivalents), and DPDS (40 equivalents) in DMSO and TEA for 6 hours, as described above. The product was purified by preparative scale HPLC on a C18 reverse phase column over 27 CVs of 0-10% acetonitrile in 25 mM TEAB buffer (pH 7.5).

### Stopped-flow measurements of hybridization kinetics

Association and dissociation rate constants for pairs of interacting oligonucleotides were measured using a Jasco FP-8500 Spectrofluorometer equipped with an SFS-852T Stopped-Flow accessory. All experiments were performed by acquiring the time-dependent emission of a 2Ap-containing strand after the addition of the complementary oligonucleotide at concentrations typically ranging from ≈ 0.5 μM to ≈ 5 μM. To analyze the data, the fluorescent traces were normalized for the signal acquired in a control experiment performed with the fluorescent oligonucleotide mixed with buffer only. To extract bimolecular kinetic constants, every experiment was independently fit in MATLAB as a bimolecular reaction, modeled using a set of differential equations solved with the variable-step solver *ODE15s* (see Supplementary Data 1, Supplementary Eq. 1 and Supplementary Figure 1A for more details).

### Analytical model for hybridization kinetics

In this work, the annealing process between two oligonucleotides A and B to form the A:B duplex is modeled following the nucleation-zippering framework (30, 31), which assumes that the hybridization process is controlled by the formation of a nucleation region composed of a few base-pairs between the two oligonucleotides, followed by the rapid zippering of the two strands. To capture the effect of pairing defects on this process, we used a variation of the simple analytical model that has recently been derived and employed (20, 32) for the study of nucleation-limited hybridization in short DNA and RNA oligonucleotides.

Briefly, this model treats the formation of any nucleation region of size *n* between oligonucleotide A (of length *L*_*A*_) and oligonucleotide B (of length *L*_*B*_) as occurring with a fixed bimolecular rate *k*_*nuc*_. Since the nucleation region is extremely unstable, its formation is quickly followed by either dissociation or zippering up to form the stable double helix. The association rate following a nucleation event in the (*i*^*th*^,*j*^*th*^) site can thus be treated as the product of the constant limiting nucleation rate *k*_*nuc*_ times the probability of successfully zippering before dissociating (*p*_i,j_^*zippering*^), leading to the following expression for the overall association rate:

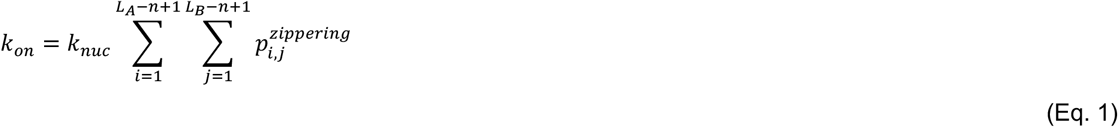

with *i* and *j* being the indexes of the first out of the *n* nucleobases constituting the nucleation site, counted in the 5′ → 3′ direction on each strand. Finally, the probability of successfully zippering after the nucleation event can be defined as the competition between (i) the two strands zippering up with a nucleation region-independent rate (*k*_*nuc → zip*_) and (ii) the two strands dissociating with a nucleation region-dependent rate (*k*^*i,j*^ _*nuc → diss*_). To capture the effect of more tightly binding nucleation regions in reducing the dissociation rate of the nucleated stretch, we can treat the nucleation region-dependent dissociation rate as equal to the product of a basal dissociation rate *k′*_*nuc → diss*_ and the nucleation region dissociation equilibrium constant 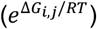, so that for *ΔG*_*i,j*_ = 0, *k*^*i,j*^ _*nuc → diss*_ reduces to *k′*_*nuc → diss*_, and for every *i*^*th*^,*j*^*th*^ nucleating region we can express *p*_*i,j*_^*zippering*^ as follows:

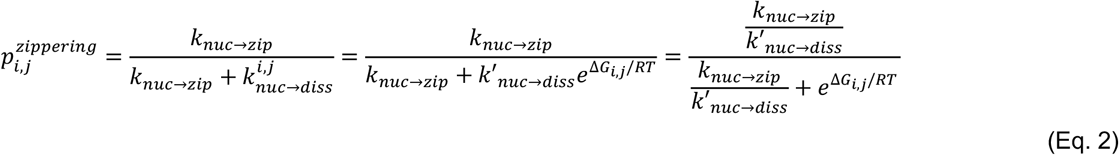

Since we are mainly interested in capturing the effect of mismatches and bulges on the probability of successfully zippering, we can rewrite 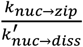 as equal to *α*, so that the overall equation becomes:

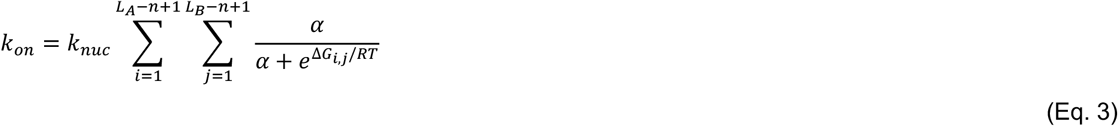

The nucleation free energy *ΔG*_*i,j*_ for non-complementary segments was fixed as equal to infinity, while for complementary segments it was computed using Nupack 4.0 at 25°C and 1M NaCl, so that the model appropriately scales the probability of detaching after nucleation according to the nucleation region binding strength. It is noteworthy that adding the length correction originally proposed in the implementation by Hertel and colleagues (32) reduces the model performance for our data, so it has been removed here. More details on the fitting procedure can be found in Supplementary Data 2, Supplementary Eq. 2 and Supplementary Figure 2. The set of estimated parameters and confidence intervals is reported in Supplementary Table 2.

### Numerical simulations for hybridization kinetics

To explicitly account for the individual steps in the zippering process, we employed a double-ended zippering model that treats hybridization as a stochastic process over the Markov-chain of in-register bound states among two strands, similarly to that shown by Schaeffer, Flamm and Mensen (33–35). The propensity for every transition between two states differing by the addition or removal of one base-pair at the ends of the zippering duplex was computed using the Metropolis method (36). Simulations were performed by fixing the unimolecular rate for the formation of a new base-pair at the rate of 4.2 × 10^8^ s^-1^ (33, 37). The hybridization rate at 1 µM of strand concentration was determined from the mean first passage time to complete zippering or detachment as implemented in the “first-step” method of Multistrand (20, 33).

### Measurement of oligonucleotide pairing thermodynamics

The dissociation constant at 25°C (*K*_*D*_, see Table 1 for list of names and definitions) for all oligonucleotide pairs studied in this work was determined by measuring the interaction of a probe sequence with a 2-aminopurine-containing target or competitor. The extent of binding was determined by normalizing to the fluorescence emission of 2-aminopurine, which is maximal when the target is in a single stranded-state and minimal when it is duplexed.

**Table 1.** Table of names.

| Name | Definition |
| --- | --- |
| $k_{on}$ | Bimolecular association rate of two oligonucleotides. |
| $k_{off}$ | Unimolecular dissociation rate of two oligonucleotides. |
| $K_D$ | Equilibrium dissociation binding constant of a duplex. |
| $k_{nuc}$ | Bimolecular rate of formation of a nucleation region between two hybridizing oligonucleotides. |
| $k_{nuc \rightarrow zip}$ | Unimolecular rate to zipper up a duplex from a nucleated state. |
| $k'_{nuc \rightarrow diss}$ | Basal unimolecular rate for the dissociation of a nucleated state. |
| $\alpha$ | Ratio between the zipping rate of the duplex and the dissociation rate of the nucleation region following its formation. |
| $R^2$ | Coefficient of determination for predicted vs measured values |
| RMSE | Root mean squared error for predicted ($\hat{y}$) vs measured values ($y$). The value is given as a percentage, and it is computed over a dataset of size $m$ as $RMSE = \frac{100}{m} \sum_{i=1}^m \sqrt{(\hat{y}_i - y_i)^2} / y_i$ |

For weakly binding oligonucleotides (*K*_*D*_ ≳ 10^−6^ M), the *K*_*D*_ was measured by direct titration of the probe oligonucleotide. For tightly binding oligonucleotides, the binding constant at 25°C could not be directly assessed, and it had to be extrapolated by studying the melting of several solutions at different oligonucleotide concentrations measured with a Jasco FP-8500 Spectrofluorometer equipped with an ETC-815 Peltier temperature-controlled cell holder. For each pair of oligonucleotides, we acquired the fluorescence emission at 370 nm while exciting at 305 nm and increasing the temperature with a gradient of 3°C/min. Every trace was independently fitted to obtain the *K*_*D*_ values at 25°C, as described previously (20). Typical results are depicted in Supplementary Figure 1B. From these values, the mean and standard deviation of the dissociation constants for each duplex were determined from a lognormal distribution. With this approach we found a modest 2Ap destabilization in DNA of +0.9 kcal/mol at room temperature, comparable with literature data (38), and a statistically non-significant destabilization in RNA at room temperature, in agreement with previous results (20, 39).

### Measurement of competition between oligonucleotides

To study competition for annealing in mixtures of oligonucleotides, we pre-mixed in buffer our target strand (T) bearing 2-aminopurine and a competitor strand (C) that differs in sequence from T only by one or two mutations. These oligonucleotides were present at twice their final desired concentrations in buffer. An equal volume of the probe sequence (P), which is fully complementary to T, dissolved in buffer at twice its final desired concentration, was then added to initiate the reaction. In this mixture, both the perfectly paired duplex T:P and the mismatched duplex C:P can be formed.

To measure the extent of hybridization of T:P, we recorded the time-dependent fluorescent signal (*F*_*mix*_) of these mixtures. The signal was normalized to yield the fraction of bound target as follows:

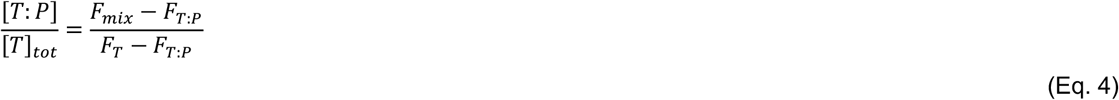

Where *F*_*mix*_ is the measured fluorescent signal coming from the mixture, *F*_*T:P*_ is the signal coming from a positive control solution containing the pre-annealed target and probe perfect duplex, and *F*_*T*_ is the signal coming from the negative control solution with the target sequence only. To accurately determine the bound fraction at equilibrium, we heated the mixed samples at 90°C for 3 minutes and then slowly cooled them to room temperature before repeating the measurements. Considering that the *K*_*D*_ of T:P in this study is significantly lower than the concentrations of the oligonucleotides employed here, it follows that all unhybridized target strands in solution must be effectively outcompeted by the competitor strand. Consequently, it has to be true that *[T:P]/([T:P]+[C:P]) = [T:P]/[T]*_*tot*_. This study allowed us to independently confirm the previously extrapolated 2Ap destabilization in DNA but not RNA at room temperature.

### Kinetics of non-enzymatic primer extension and ligation in competing mixtures

Since primer extension reactions require Mg^2+^ to proceed, we used a different buffer containing 200 mM Tris pH 8.0 and 100 mM MgCl_2_, that does not to significantly alter the overall equilibration process (20). All primer extension and ligation reactions were performed at 10-µL scale, with 1 µM probe sequence, 1 µM perfectly matched target sequence (5′-FAM labeled), 1 µM mismatched competitor sequence (5′-Cy5 labeled), 200 mM Tris (pH 8.0), and 100 mM MgCl_2_. For each reaction, the probe was first mixed with Tris 8.0 and MgCl_2_ at the bottom of the tube. Then, target and competitor were loaded separately on the lid or wall of the tube. The stock solution of C*C/*CGCA was prepared freshly and added into the tube, followed by immediate spin down and mixing to initiate the reaction.

To sample the reaction at different time points, 0.5 µL of the reaction mixture was transferred to 25 µL of a quenching buffer composed of 25 mM EDTA, 1X TBE, and 2 µM RNA strands complementary to the probe, in formamide to dissociate the dye-labeled strands from the probe. The products of target and competitor reactions were resolved by 20% (19:1) denaturing PAGE and the images in the green and red channels were acquired using a Typhoon 9410 scanner to detect both FAM-labeled and Cy5-labeled oligos. The reaction yield over time was quantified using ImageQuant TL software.

## RESULTS

### Sequence design

To characterize the effect of defects on the annealing behavior of DNA and RNA, we designed a target (T) oligonucleotide 5′-TGGTGATGCGTG-3′ and systematically edited its sequence introducing mismatches or bulges to produce competitors (C). All these sequences were tested for binding to a probe (P) of fixed sequence 5′-CACGCATCACCA-3′. Eevery substitution in the competitors was picked to minimize secondary structures and self-pairings as predicted by Nupack (15).

All target sequences and substitutions used for kinetic and thermodynamic characterization are listed in Supplementary Table 1, with every sequence name referencing the nature and the position of the defects in the 5′ → 3′ direction, either mismatches (m) or bulges (b), along the double helix. For example, the sequence b_4_ has a deletion that introduces a bulge in position 4 of the probe, and m_3_ introduces an internal mismatch in position 3 of the probe. Whenever an RNA sequence is used in this work, all deoxyribonucleotides are substituted by their respective ribonucleotides, and every deoxythymidine is substituted by a ribouridine. Sequences used to study the kinetics of non-enzymatic primer extension and ligation in competing mixtures are listed in Supplementary Table 3.

### Effect of defects on short oligonucleotide hybridization

When two fully complementary target (T) and probe (P) oligonucleotides are mixed in solution, they quickly hybridize with a second order rate constant *k*_*on*_ to produce a double helix T:P. In this work we investigated the effect of mismatches and bulges on hybridization by systematically altering the position and nature of such defects, yielding the results shown in Figure 1A. We found that the mismatches and bulges we studied did not affect the *k*_*on*_ values by more than one order of magnitude with respect to the control sequence, which forms a perfect 12 bp-long duplex (t12). As expected, most defects do slightly reduce the rate of formation of the duplex, with association rates for the 12-mer DNA oligonucleotide affected the most by double mismatches, followed by single mismatches and least by bulges.

**Figure 1.**
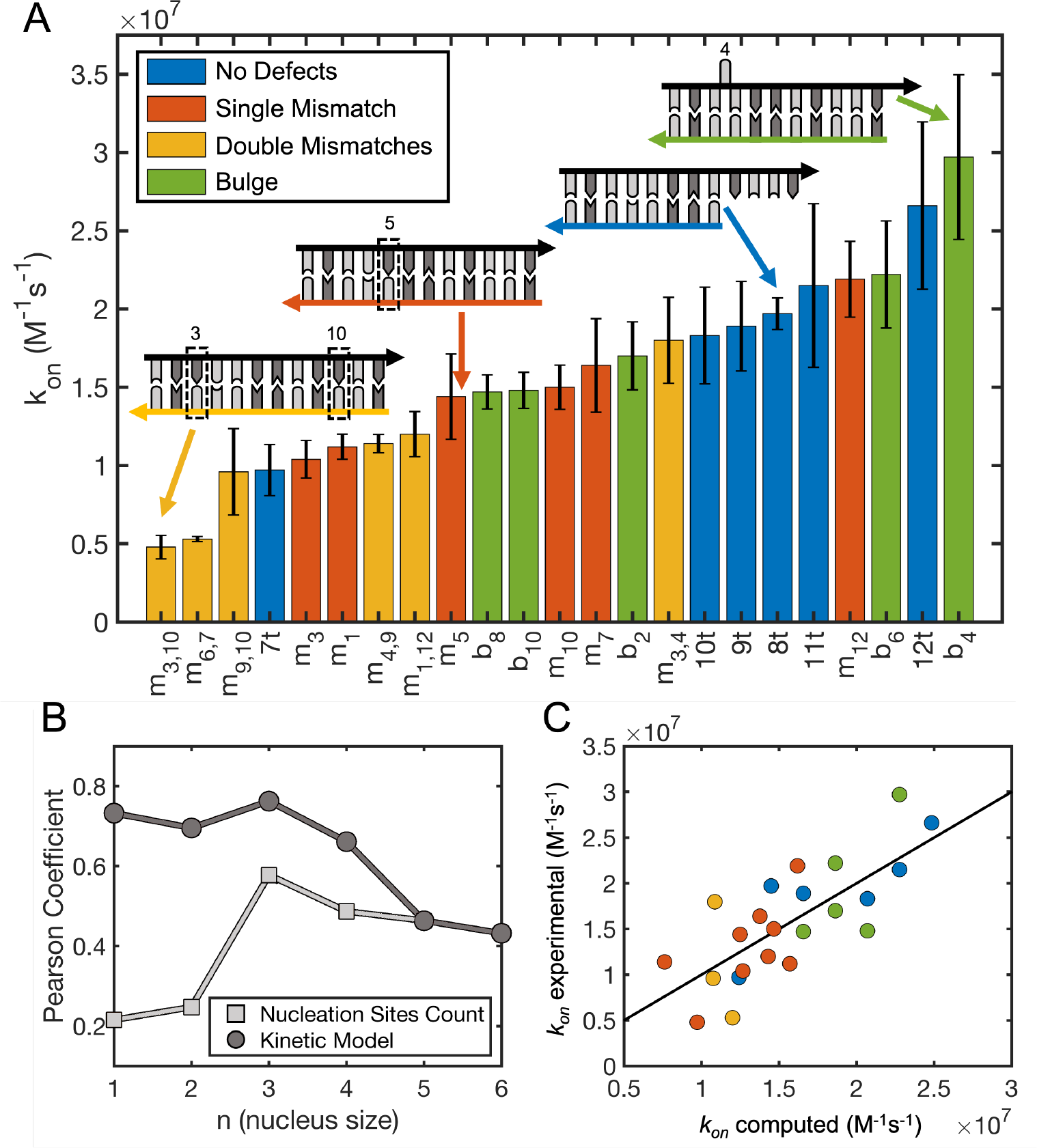
Hybridization kinetics for short DNA oligonucleotides with defects. (**A**) Hybridization kinetics constants for short DNA oligonucleotides with and without single mismatches, double mismatches, or bulges. Data have been sorted from lowest to highest hybridization rate to facilitate comparisons with the control (12t), which forms a perfectly pairing 12 bp-long duplex. The sketches show the general sequence designs, with the probe sequence (in black) fixed while changing the pairing oligonucleotides. (**B**) Correlation between experimentally determined hybridization rates and (light gray) the total number of nucleation sites or (dark gray) rates predicted by the kinetic model from Eq. 1 constructed with nucleation sites of different sizes. (**C**) Hybridization rates computed with the kinetic model described in Eq. 1 using n=3 against experimentally determined rates.

By introducing defects in different positions along the duplex, we continuously reduced the length of the longest continuous stretch of complementary nucleotides down to 4. Interestingly, by doing so we did not see any clear discontinuity in the measured hybridization rates, suggesting that “the rule of seven” as formulated by Cisse and colleagues (26) does not apply to our system. In the nucleation-zippering framework, hybridization is treated as a process controlled by the formation of a nucleation region composed of a few base-pairs between the two oligonucleotides, followed by the rapid zippering of the two strands (30, 31). The nucleation rate is generally regarded as the limiting step, with a nucleation region forming only once in every ∼10^3^ collisions (20, 33). The total hybridization rate is then the sum of multiple hybridization pathways that can be initiated at different nucleation sites. In this framework, a net reduction in the total number of available nucleation sites due to the introduction of defects would inevitably reduce the total hybridization rate (20, 32). By systematically computing the number of available nucleation sites as a function of nucleation site length, we found that the best correlation with the association rates we measured was for continuous 3 nt stretches of complementarity (Pearson correlation coefficient = 0.58, Figure 1B), analogously to what was recently shown by Hertel and colleagues for perfectly pairing DNA oligonucleotides (32).

However, the moderate overall correlation masks the fact that the hybridization rates for perfectly pairing oligonucleotides and oligonucleotides with bulges are much better correlated with the total number of available nucleation sites (Pearson coefficient = 0.72). In contrast, the variation in the association rate for mismatched oligonucleotides is not well captured (Pearson coefficient = 0.50). We hypothesized that the model could not describe all hybridization rates at the same time because of a reduced zippering rate for mismatched oligonucleotides, which must propagate their zippering across non-canonical base pairings to successfully form a stable double helix, consistent with literature on the folding energy landscape of hairpins with mismatches (40, 41) and a more recent assessment on short DNA duplexes containing abasic sites (42).

To capture this effect, we employed a simple nucleation-limited analytical model that describes the hybridization phenomenon (32). In the context of this model, two randomly diffusing strands can collide and form a nucleation region composed of a few base-pairs. This transient event can be followed either by the detachment of the two strands or their rapid zippering to form the stable double helix. In this process, the main variables that control the overall outcome are the nucleation rate (*k*_*nuc*_) and the ratio (*α*) between the zippering rate of the duplex and the dissociation rate of the nucleation region. The model was fit to our data using a single shared nucleation rate and an additional defect-dependent *α*. This correction greatly improves the model performance (Pearson coefficient = 0.76, Figure 1C) and results in a best fitting nucleation rate equal to 2.2 × 10^6^ M^-1^s^-1^. Since the estimated *p* _*i,j*_ ^*zippering*^ for perfectly paired or bulged sequences is extremely close to unity with n=3 as length of the nucleation region, we could only reliably estimate lower boundaries for *α* as equal to 7.2 × 10^−4^ and 5.4 × 10^−4^ respectively. In contrast, values of *α* for mismatched and double mismatched sequences are well constrained, with best fitting values equal to 4.3 × 10^−5^ and 4.6 × 10^−5^ respectively, leading to a median *p* _*i,j*_ ^*zippering*^ equal to ∼ 0.7 over all possible nucleation sites available for n=3. With this study, we found that *α* for mismatched sequences is significantly smaller than for perfectly paired or bulged sequences, reflecting a significantly lower success rate in zippering after nucleation region formation for single and double mismatches but not for bulges. It is important to emphasize that this simplified model does not explicitly consider each individual step of the zippering process. To obtain a more physically grounded insight into the hybridization process, we implemented a numerical simulation, as described in the Materials and Methods, that explicitly accounts for the whole zippering trajectory starting from any possible nucleation region composed of a single base-pair between in-register complementary nucleotides. Assuming that the presence of mismatches in the sequence only affects the process of zippering but not the rate of formation of the initial nucleus, we found a best fitting nucleation rate equal to 2.0 × 10^6^ M^-1^ s^-1^, and an overall best fitting energy barrier to overcome internal mismatches in the zippering process equal to 2.49 kcal/mol. In the context of this model, mismatches reduce the hybridization association kinetics by causing a drop in the success rate when the zippering starts from a position adjacent to a defect and by dramatically increasing the time required to complete the hybridization. According to both approaches, bulges do not interfere with downstream zippering of the duplex during hybridization.

To extend our findings to RNA oligonucleotides, we studied the RNA counterparts of a subset of the 12-nt DNA sequences discussed here. In this case, defects also cause the association rate to vary, again within only one order of magnitude from the control. The impact of the different types of defects was similar to that measured in DNA, with a small if any detectable effect due to a bulge or single mismatch and a large effect due to double mismatches. As a biologically relevant test case, we measured the effect of a single mismatch in the 8 bp long miR125 seed sequence, and found that the presence of the defect does not significantly affect the hybridization on rate (Supplementary Data 3 and Supplementary Figures 3 and 4). This result extends our findings to shorter sequences and lower ionic strength, and is in direct contrast to what has previously been reported (26). This suggests that our finding of a minor effect of defects on association kinetics for 12 bp-long duplexes may apply more generally to shorter and longer sequences.

### Effect of mismatches and bulges on the off-rates of short oligonucleotides

The eventual separation of two hybridized strands occurs after a characteristic time described by the first-order rate constant *k*_*off*_. To determine the *k*_*off*_ for our duplexes, we employed the relationship between the equilibrium dissociation constant *K*_*D*_, and the association and dissociation rate constants, so that under the assumption of a two-state transition:

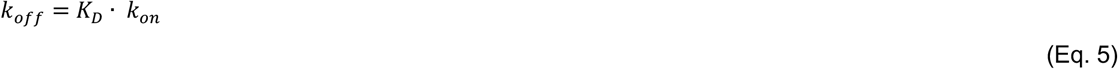

We determined *K*_*D*_ at room temperature for our duplexes through a series of melting and binding experiments. Given *K*_*D*_ and *k*_*on*_, we could calculate *k*_*off*_, thus estimating the rate at which the strands of a duplex dissociate. The values of *k*_*on*_ and *k*_*off*_ for DNA determined in this work are shown in Figure 2 on a logarithmic scale to better emphasize the relative contribution of forward and reverse reactions to defect-mediated destabilization. Our data show that the main overall contribution of defects to duplex destabilization is on the dissociation rate constant. For all defects studied here, we find that the destabilization is both position-dependent and consistent with NN predictions (see Supplementary Data 4 and Supplementary Figure 4). We confirmed prior observations (20, 42–44) that the logarithm of the dissociation rate is well-correlated with the free energy of pairing of two oligonucleotides both in DNA and RNA (R^2^ = 0.97, Supplementary Figure 3D), even when defects are present. By treating dissociation as a thermally activated process (43–45), we can compute an average reduction in the activation energy caused by the defects in a DNA duplex as equal to -5.0 kcal/mol, -3.8 kcal/mol, and -2.6 kcal/mol for double mismatches, bulges, and single mismatches respectively, following a trend that is different from that observed for *k*_*on*_, where bulges do not significantly affect the process.

**Figure 2.**
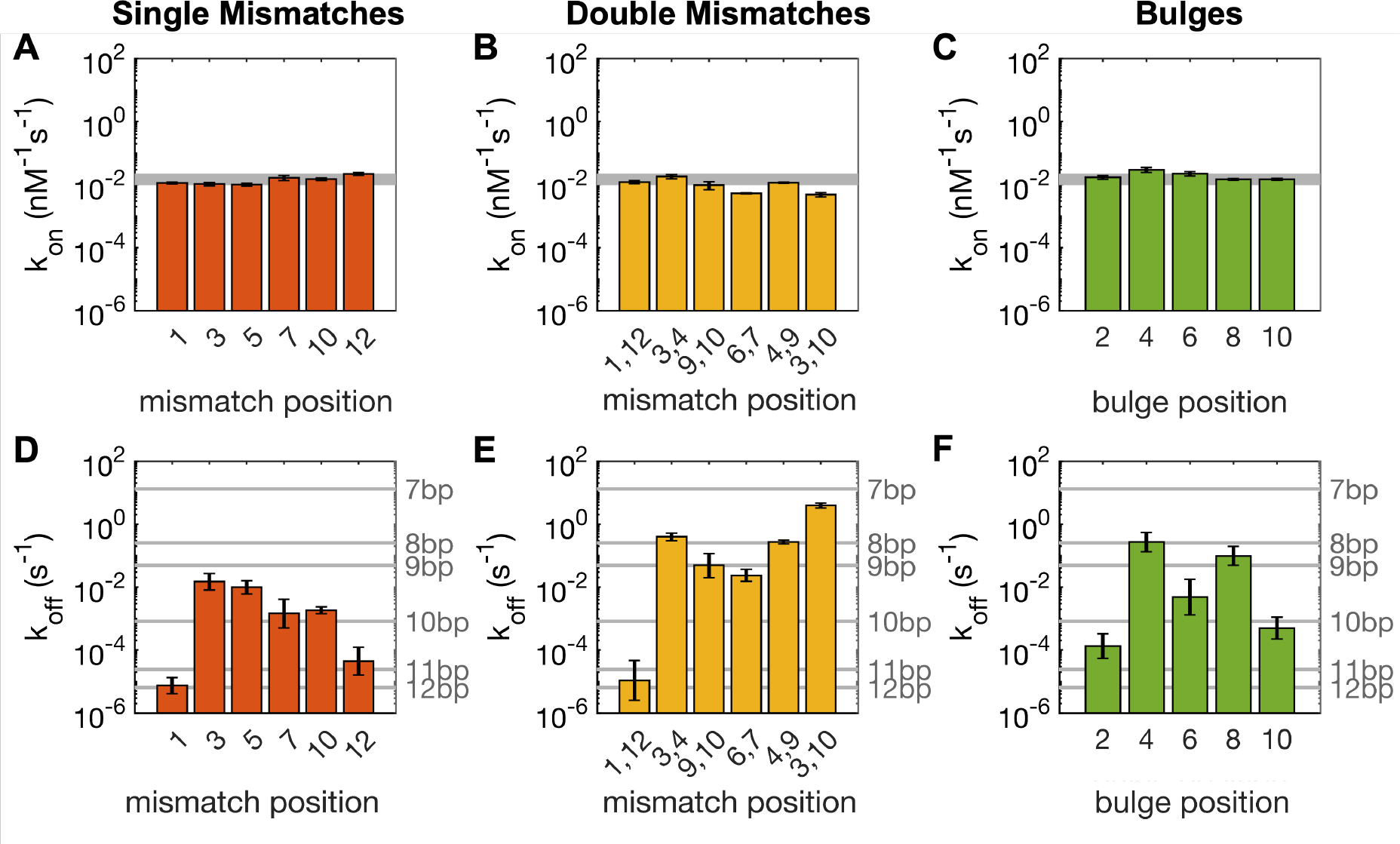
Kinetics for association and dissociation in short DNA oligonucleotides with defects. (**A, B, C**) Association rate constants for oligonucleotides with either single mismatches (**A**), double mismatches (**B**), or bulges (**C**). The grey shaded bands show *k*_*on*_ for control oligonucleotides of lengths varying from 7bp to 12bp, yielding values varying by roughly a factor of two. Data are shown on a logarithmic scale to directly compare the relative contributions of *k*_*on*_ and *k*_*off*_ to *K*_*D*_. (**D, E, F**) Dissociation rate constants for short DNA oligonucleotides with either single mismatches (**D**), double mismatches (**E**), or bulges (**F**). Horizontal lines show *k*_*off*_ for a series of perfectly paired control duplexes.

### Competition for binding of short oligonucleotides

We next extended our investigation to the competition between defect-containing oligonucleotides and a perfectly complementary oligonucleotide for binding to a common probe oligonucleotide. When we add a target oligonucleotide (T) designed to bind to a perfectly complementary probe oligonucleotide (P) in a complex mixture in the presence of another competing strand sharing a partial complementarity (C) to the probe, we expect a certain fraction of both T:P and C:P duplexes to form initially (Figure 3A). In this scenario, every C:P duplex can be considered to be an off-target species. To assess the extent of target binding at any time, we monitored the emission of a 2Ap nucleotide within the target sequence as outlined in the Materials and Methods section. In order to quantify the interference caused by the competitor, we introduced a parameter termed binding accuracy (*θ*), defined as the ratio of correctly bound probe to the total bound probe:

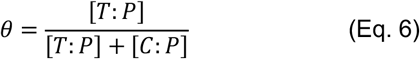

**Figure 3.**
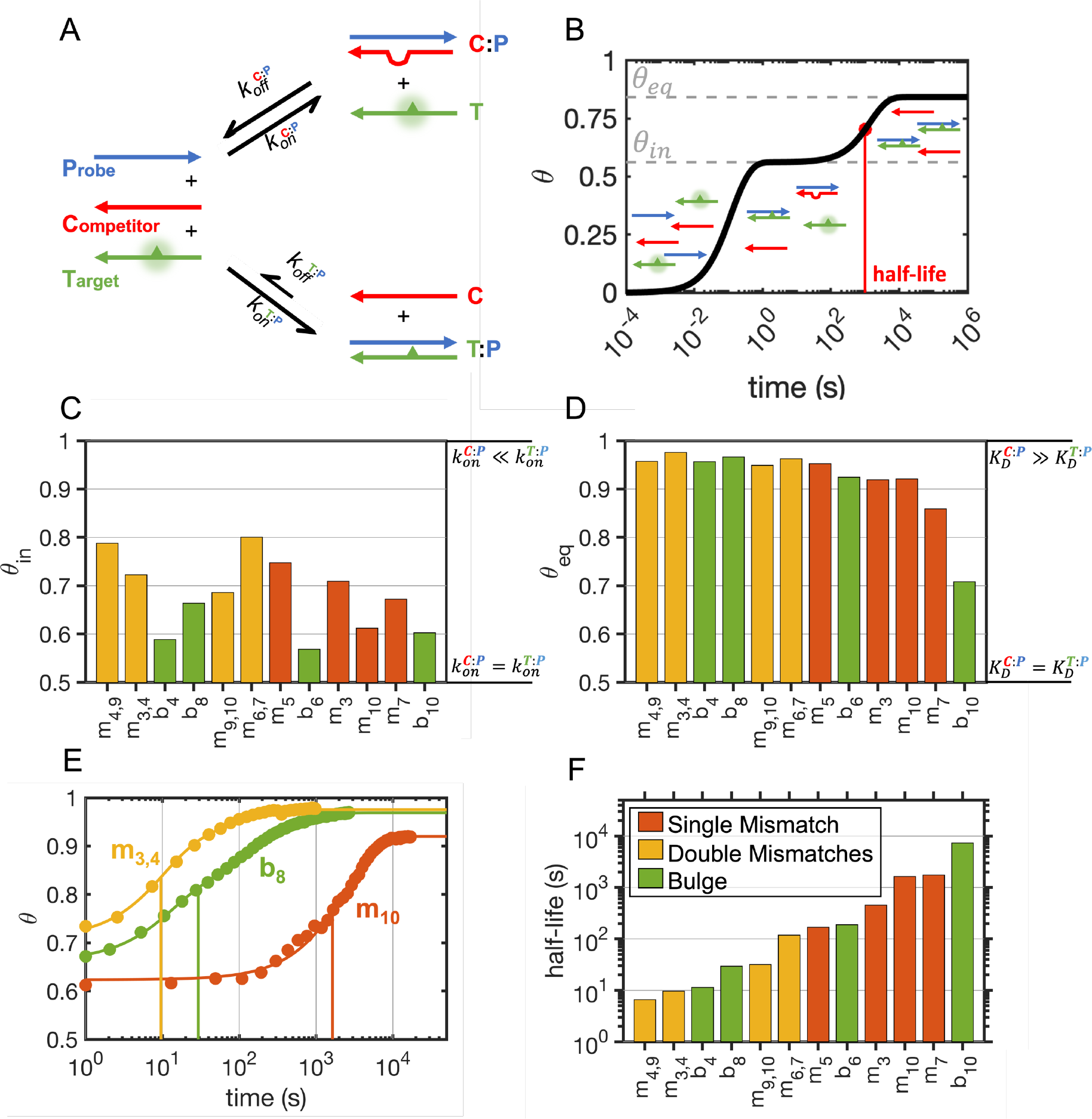
Competition experiment for DNA probe hybridization. (**A**) Reaction scheme for the competition experiment between equimolar amounts of probe sequence (P), perfectly pairing target (T) and defected competitor (C). Target strands bear a fluorescent nucleotide (2Ap), sketched as a triangle, which allows tracking of the bound fraction over time. (**B**) Sketch of the time course of a typical competition experiment: the three strands are mixed and hybridize to produce a certain fraction of perfect and imperfect duplexes in ≈ 1 second, so that the system temporarily exhibits the initial binding accuracy *θ*_*in*_. After a certain amount of time depending on *k*_*on*_, *k*_*off*_ and strand displacement rates, the metastable state converges to the energy minimum that is associated with its equilibrium binding accuracy *θ*_*eq*_. (**C**) Initial binding accuracy for competition experiments in case studies where *θ*_*in*_ < *θ*_*eq*_. These values are expected to be equal to 1.0 for 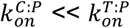 0.5 for 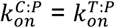 (**D**) Binding accuracy at equilibrium for competition experiments in case studies where *θ*_*in*_ < *θ*_*eq*_. These values are expected to be equal to 1.0 for 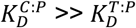 or 0.5 for 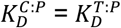 (**E**) Equilibration traces measured in competition experiments. The continuous lines were derived by interpolating/extrapolating the data. Vertical lines highlight half-lives for the shown samples. (**F**) Measured equilibration timescale, defined as the time required so that 50% of the transient state is equilibrated (see panel B). The transient low-accuracy state can last for a time ranging from a few seconds up to hours. All experiments were performed using 200 nM of each strand in 10 mM Tris-HCl pH 8.0 and 1 M NaCl.

Drawing upon our understanding of the behavior of complex mixtures of competing oligonucleotides (20), we expect rapid formation of both C:P and T:P duplexes in ∼1 second. If the dissociation rates of the duplexes are significantly slower than their initial rates of formation, the relative amount of the two duplexes initially formed upon mixing of the three strands will depend uniquely on their relative association rates (Figure 3B). In this early stage, an initial binding accuracy (*θ*_*in*_) could be conveniently measured, typically assessed after approximately 2 seconds from hand mixing. Following the initial binding event and the partitioning of the probe oligonucleotide between perfectly (T) and imperfectly (C) complementary pairing partners, the system progressively evolves and converges to an equilibrium state characterized by a stable binding accuracy, defined here as *θ*_*eq*_.

In most cases studied here using the 2Ap-containing oligonucleotide 12t as the target sequence, we found that the initial binding accuracy (Figure 3C) differs from the equilibrium binding accuracy (figure 3D), with *θ*_*in*_ < *θ*_*eq*_. Thus, as the system approaches equilibrium, we see an increase in hybridization accuracy (*θ*). The only exceptions to this were those cases where the presence of 2Ap caused a small increase in *K*_*D*_ that is comparable to the increases caused by variations in the competitor strand, namely m_1_, m_12_, m_1,12_ where we could measure a decrease in *θ* as a function of time, and b_2_, where we see that *θ*_*in*_ = *θ*_*eq*_.

Having established the presence of transient low-accuracy states in a competitive scenario, our investigation shifted to understanding the significance of such states. Specifically, we aimed to determine whether these mismatched duplex states are short-lived or long-lived relative to the timescales of processes such as nonenzymatic template copying where high specificity is required in order to minimize the propagation of errors. To this end, we conducted a comprehensive study of the evolution of the 2Ap signal under conditions where the binding accuracy increases over time. Equilibration within a multi-strand mixture is a complex process that may not follow a single exponential decay (20), making it challenging to define a simple equilibration rate (Figure 3E). Consequently, we chose to define the timescale for equilibration as the time required for 50% of the transient imperfect duplexes to disappear (Figure 3B,E). We found that the timescales associated with the equilibration process vary by several orders of magnitude, with sequences bearing double mismatches taking only a few seconds to equilibrate in the fastest cases, while sequences with single mismatches or bulges taking hours in the slowest cases (Figure 3F).

### Prediction of initial binding accuracy for short oligonucleotides

Having established that imperfect duplexes can be transiently present for a relatively long time as the system equilibrates, we wanted to know whether their initial abundance could be predicted. Taking into account the modest destabilizing factor coming from 2Ap (+0.9 kcal/mol), we could calculate the exact analytical solution for the expected *θ*_*eq*_ (46) by knowing the total concentration of the three strands and the binding constants (*K*_*D*_) for each duplex. Moreover, we could conveniently approximate predictions for the expected *θ*_*in*_ as follows for equimolar concentrations of C and T strands:

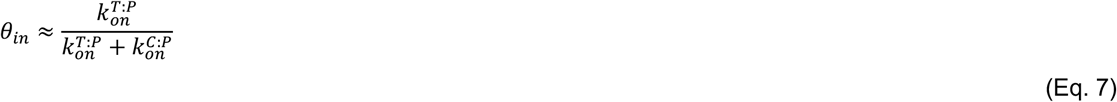

It is important to note that Equation 7 reduces to *θ*_*in*_ = 1.0 in the case where 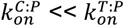 Similar association rates are required to have *θ*_*in*_ ≠ *θ*_*eq*_ in those cases when the target is designed to outcompete the competitor at equilibrium.

We found that both the measured initial (R^2^ = 0.60, RMSE = 8.8%) and equilibrium (R^2^ = 0.89, RMSE = 6.5%) binding accuracies are moderately well predicted from the known association and dissociation rates (Figure 4A,B). Importantly, *θ*_*in*_ and *θ*_*eq*_ are poorly correlated (R^2^ = 0.19, RMSE = 32%) as depicted in Figure 4C, showing that knowledge of the equilibrium state of the system is of little value when considering the out-of-equilibrium regime.

**Figure 4.**
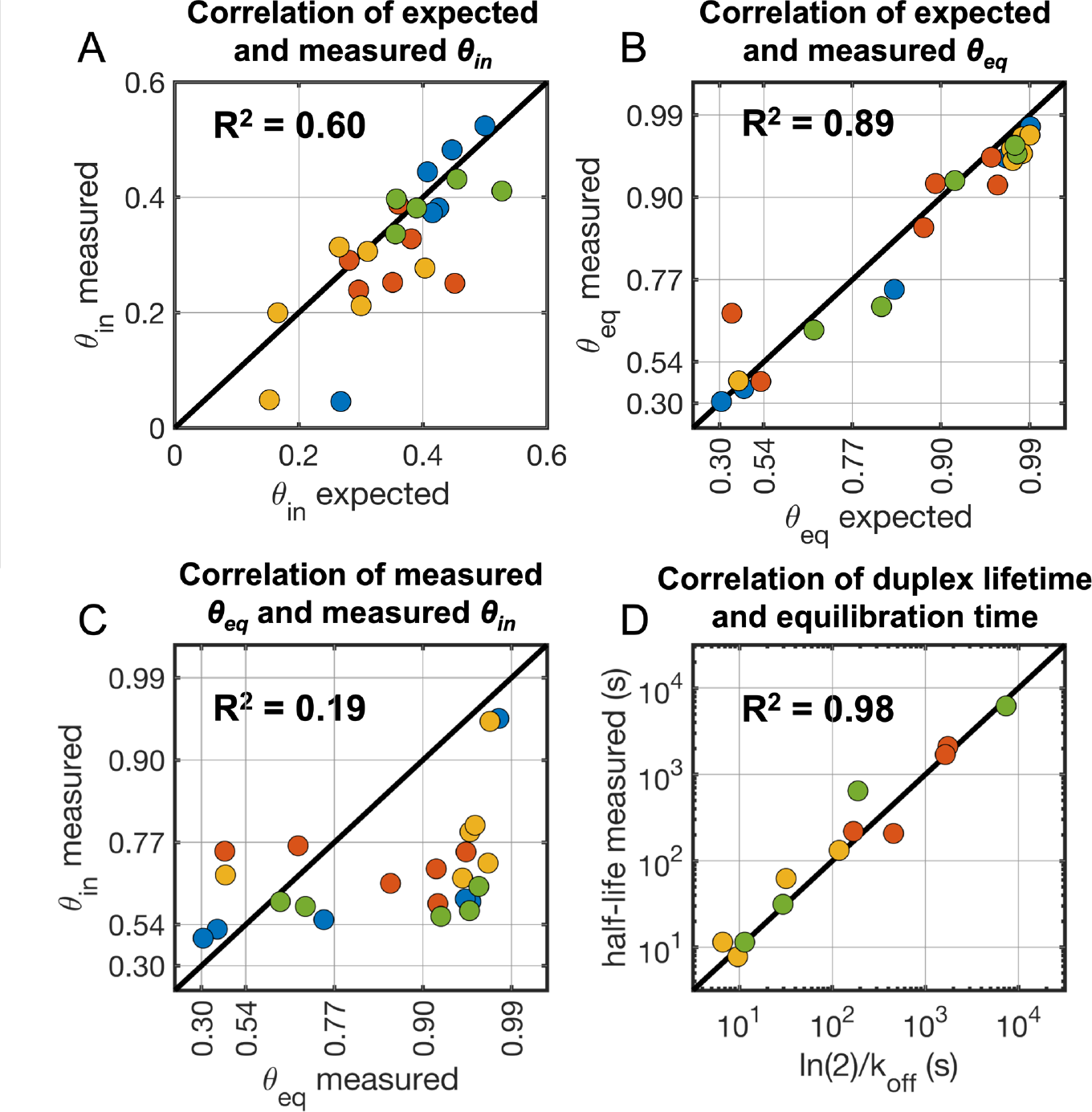
Predictions for competition experiments with DNA. (**A**) Initial binding accuracy (*θ*_*in*_) for the competitive mixtures compared to values computed from Eq. 7 are in good agreement, showing that association rates can be used to predict the initial partition of duplexes. (**B**) Binding accuracy at equilibrium is well predicted by a simple binding model based on experimentally determined nucleic acids dissociation constants. Axes have been stretched to better visualize the cluster of points close to 1.0 (**C**) Partition at equilibrium and after initial mixing are not well-correlated, showing the importance of expanding our current models for complex mixtures of oligonucleotides. Red dashed line shows best fitting line. Axes have been stretched to better visualize the cluster of points close to 1.0. (**D**) Comparison of the measured equilibration timescale and the lifetime of the defect containing duplex as computed from the dissociation rate of the competitor oligonucleotide. All experiments were performed with 200 nM of each strand in 10 mM Tris-HCl pH 8.0 and 1 M NaCl.

Finally, we observed a strong relationship (R^2^ = 0.98) between the dissociation rates of the imperfect duplexes and the duration of the equilibration process. This observation suggests that there is a negligible role for strand displacement in the evolution of the system at these concentrations. This outcome is a natural consequence of the intentional absence of toeholds in the design of the imperfect duplexes, resulting in very slow strand displacement (47) so that the overall equilibration process is controlled by the dissociation and reshuffling of the strands.

Interestingly, we found that this is not the case for competition among the RNA strands (Supplementary Data 6 and Supplementary Figure 6) studied in this work: their much slower dissociation rates compared to DNA (20, 43) would result in years-long transient states that do not match our experimental results. This discrepancy can be understood in the context of a major role for strand displacement: given the measured lower boundary for the strand displacement rate of ≈ 1 M^-1^s^-1^ for RNA in the absence of toeholds (20, 47) and the major enhancement caused by the presence of mismatches (48, 49), we can reasonably expect an upper limit of several hours for the equilibration in a competition scenario with 1 µM oligonucleotides. Thus, strand reshuffling in RNA is effectively dominated by strand displacement. Still, the stronger pairing between RNA strands makes the equilibration up to 30 times slower than for the corresponding DNAs. Thus out-of-equilibrium imperfect RNA duplexes can be abundant and long-lasting species that are potential substrates for chemical and enzymatic reactions.

### Off-target reactions in competing RNA mixtures

The fidelity of nonenzymatic template copying reactions could be greatly affected if a mismatched primer anneals to a template and becomes extended before equilibration leads to replacement of the mismatched primer with a fully complementary primer. To test whether the imperfect duplexes present in an out-of-equilibrium RNA mixture can behave as substrates for primer extension and ligation reactions, we studied a mixture of RNAs in which perfectly pairing target and mismatched competitors compete for binding to a longer probe oligonucleotide. We then examined the extension of either target or competitor using the highly reactive imidazolium-bridged C*C dinucleotide (50), and target or competitor oligonucleotide ligation with the much more slowly reacting *CGCA tetranucleotide activated with a 2-aminoimidazole moiety on its 5′ terminus (51). The imidazolium-bridged C*C dinucleotide is known to chemically extend the 3′-terminus of a paired oligonucleotide over a typical timescale of minutes at saturating concentrations (29), and over a timescale of hours at lower concentrations. The activated tetranucleotide is known to ligate to the 3′ end of a paired oligonucleotide over a timescale of days at saturating concentrations (51). These prebiotically plausible reactions are the two classes of reactions that are thought to have contributed to nonenzymatic RNA replication prior to the evolution of ribozyme polymerases in the RNA World (6, 52).

To monitor the extent of chemical primer extension and ligation, the competing oligonucleotide C (bearing the double mismatch) was labelled with a 5′-Cy5 fluorescent dye, while the target oligonucleotide T was labelled with a 5′-FAM fluorescent dye. Denaturing polyacrylamide gels were imaged in the green and red channels to quantify the reacted target strand and the reacted competitor strand over time (Figure 5B,C and Supplementary Figure 7).

**Figure 5.**
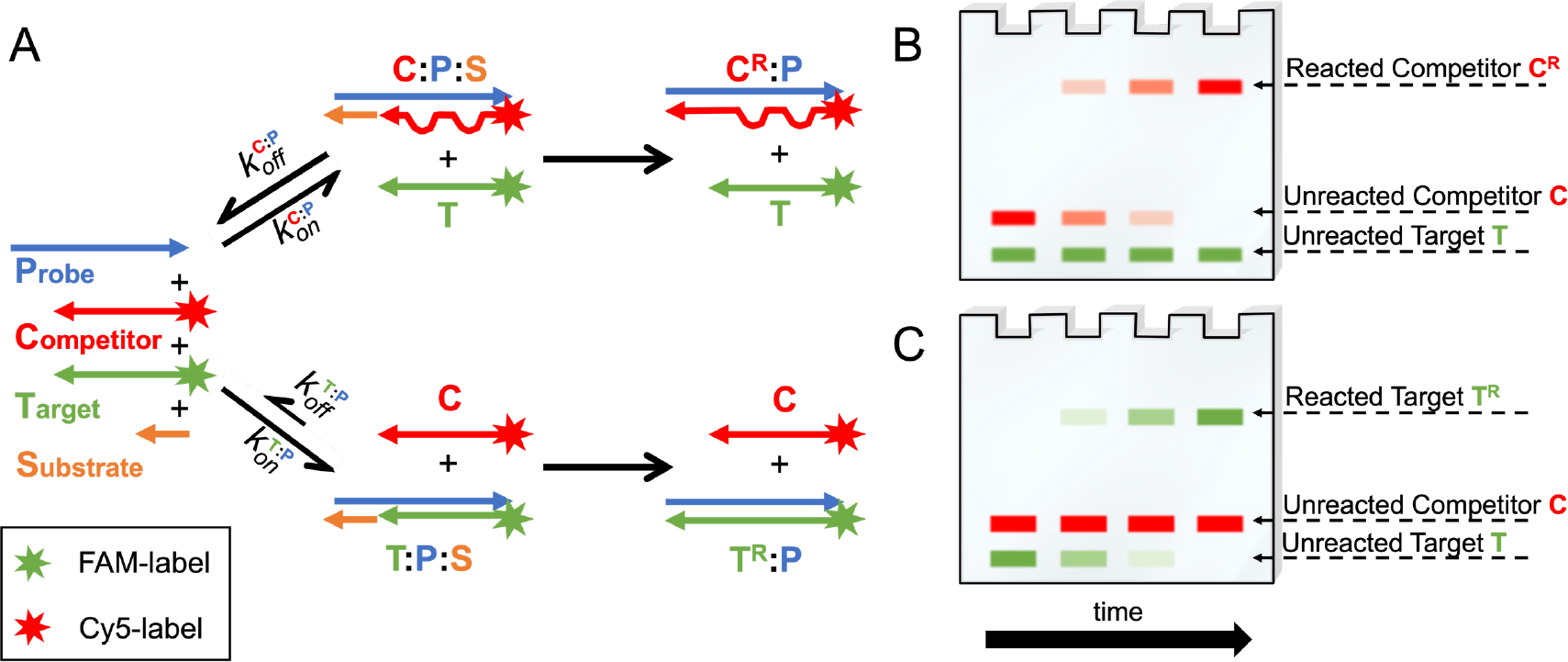
Experimental design for primer extension and ligation reactions in a competition experiment. (**A**) Reaction scheme for the competition experiment between equimolar amounts of probe sequence (P), perfectly paired target (T) and mismatched competitor (C) in the presence of chemically reactive substrates (S). Both C:P and T:P duplexes can react with the substrates to form longer species C^R^ and T^R^. The probe oligonucleotide presents either the 5′ overhang (5′-GGGG-3′) to allow binding of imidazolium-bridged C*C dinucleotides, or a different 5′ overhang (5′-UGCG-3′) to allow binding of a tetranucleotide substrate. Primer extension or ligation follows as depicted in the sketch. (**B, C**) Sketches of denaturing PAGE to resolve reaction products in a competition experiment. Both competitor, target sequences and reaction products can be tracked and quantified at the same time due to the distinct conjugated fluorophores. In the examples presented here, we show (**B**) the hypothetical outcome of a reaction where *θ* is always equal to zero, so that only the mismatched C:P duplex is formed at any time, and (**C**) the hypothetical outcome of a reaction where *θ* is always equal to one, so that only the perfect T:P duplex is formed at any time. An example of a real PAGE overlay is shown in Supplementary Figure 7.

The equilibrium binding accuracy for the mixture with m_4,9_ is predicted to be equal to 1.00, a value in agreement with our experimentally determined value of 0.97±0.02. Under a conventional description that only accounts for the equilibrium state, virtually no C:P complex would be available as a substrate for primer extension or ligation reactions. In contrast, our results show that even in the presence of a double mismatch, the competitor can be a substrate for both non-enzymatic primer extension (Figure 6A,B) and ligation (Figure 6C) during the equilibration phase, with a strikingly large yield as high as ≈ 38%. As expected, once the metastable imperfect duplexes are removed from the pool by equilibration, the mismatched competitor can no longer be elongated by either of the non-enzymatic reactions.

**Figure 6.**
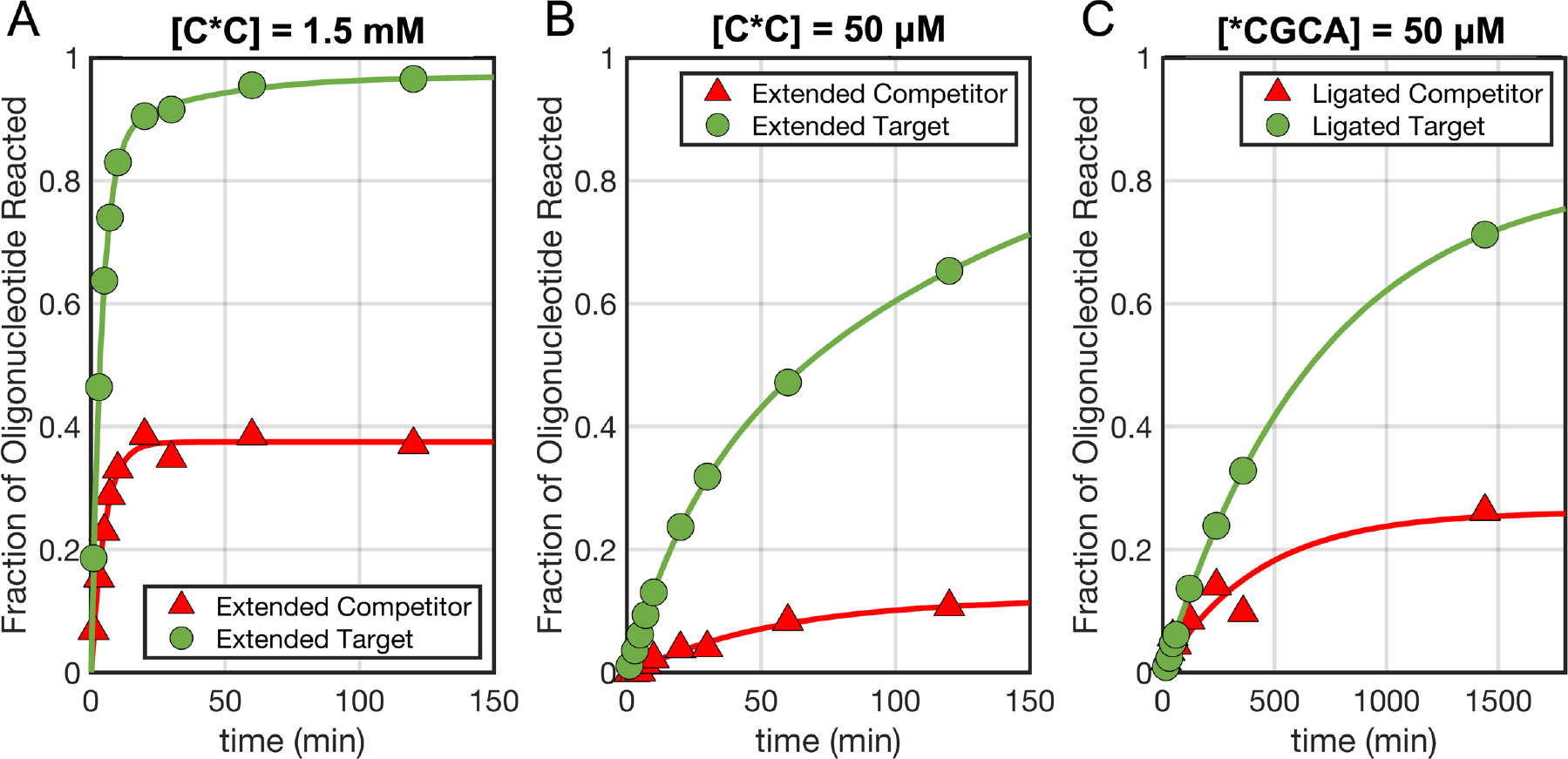
Chemical reactions in a competition scenario among RNA strands. (**A, B**) Primer extension reactions at different C*C substrate concentrations and (**C**) ligation reactions of perfect (Target) and mismatched (Competitor) strands. Continuous lines were obtained by interpolating/extrapolating the data. All reactions have been performed at room temperature in 200mM Tris-HCl pH 8.0 and 100mM MgCl_2_.

## DISCUSSION

We have studied the effect of single mismatches, double mismatches, and bulges on the association and dissociation kinetics of short DNA and RNA duplexes. We find that these defects have a negligible or minor effects on the association rate. Double mismatches affect the oligonucleotide on-rate the most, with a maximum ≈ 6x slow-down, while bulges have the smallest effect. We found that *k*_*on*_ for perfectly paired DNA sequences is well correlated with the ΔG of hybridization (R^2^ = 0.76), as previously observed for short RNA oligonucleotides (20). This relationship does not hold for imperfectly matched sequences (R^2^ = 0.08), showing that mismatches and bulges increase the complexity of the hybridization phenomenon in a non-trivial way. We then employed a simple nucleation-limited model and a kinetic Monte Carlo model to fit the experimentally determined hybridization rates, finding that the decrease in association rate can be explained in terms of loss of nucleation sites and an added energy penalty for zippering two imperfectly complementary strands. We found that our results can be qualitatively extended from DNA to RNA, and to lower ionic strength and shorter sequences.

Our study on the combinatorics of nucleation sites suggests that 3 contiguous base-pairs are necessary and sufficient to make an effective nucleation site (Figure 1B), even in the context of sequences that are not entirely complementary. No clear discontinuity in the measured hybridization rates was observed when reducing the length of the longest continuous stretch of base pairs from 12 to 4, suggesting that “the rule of seven” as formulated by Cisse and colleagues (26) may be less general than previously thought, and emphasizing the need for an orthogonal and more general study of oligonucleotide association kinetics in solution.

Our rate analysis is consistent with mismatches creating energy barriers for zippering, similarly to the effect of abasic sites on hybridization kinetics (42). The absence of this energy barrier in the zippering process over bulges is consistent with structural data showing that stacking tends to be preserved in bulged duplexes, especially when the extra base is looped-out leaving the duplex virtually unaltered (53, 54). On the other hand, mismatches cause significant structure alterations to duplex geometry which do seem to impede zippering (55). In contrast to the minimal effect of bulges on association rates, the effect of bulges on duplex dissociation is large and well-captured by the NN model of duplex thermodynamics. While this observation points to an asymmetric role of bulges in pairing and unpairing, it makes the simple thermodynamic model of duplex stability a useful tool for the prediction of dissociation kinetics for imperfect duplexes.

As a consequence of the similar association rates of perfectly and imperfectly complementary oligonucleotides, we have found that a significant fraction of imperfect duplexes form initially and exist transiently when such oligonucleotides anneal in a mixture. Even when imperfect duplexes are not expected to exist at equilibrium, off-target binding may persist over time scales that can be longer than prebiotically relevant RNA copying reactions (29, 56) or biologically relevant phenomena such as RISC-mediated RNA degradation (57, 58) or Cas9 target cleavage (59). This raises the question of the impact of such incorrect association events in biological or prebiotic contexts (6) that are far from equilibrium. To quantify the likelihood of observing out-of-equilibrium mispairings in prebiotic or biological mixtures of oligonucleotides, we have derived a simple analytical relationship (Supplementary Eq. 3 and 4) between the length of a given oligonucleotide and its probability of binding to a partially complementary strand in a mixture of random sequences (Supplementary Data 8 and Supplementary Figure 8).

Consider an out-of-equilibrium primordial environment hosting nonenzymatic RNA replication reactions and subject to a continuous influx of energy (22). Given appropriate fluctuations in temperature, salt concentration or pH, a population of oligonucleotides would have undergone a continuous change of base-paired configurations while being cyclically re-equilibrated (22). Such shuffling of annealed primer/template combinations is required for replication in our recently proposed Virtual Circular Genome model for primordial RNA replication. However, if a primer is mis-paired with an incorrect template, or if either the primer or the template contain a mutation, then primer extension or ligation will lead to propagation of incorrect information (Figure 7). It is therefore important to understand the lifetimes of such incorrect primer/template complexes, in order to understand the effects of such mispairing on the fidelity of replication.

**Figure 7.**
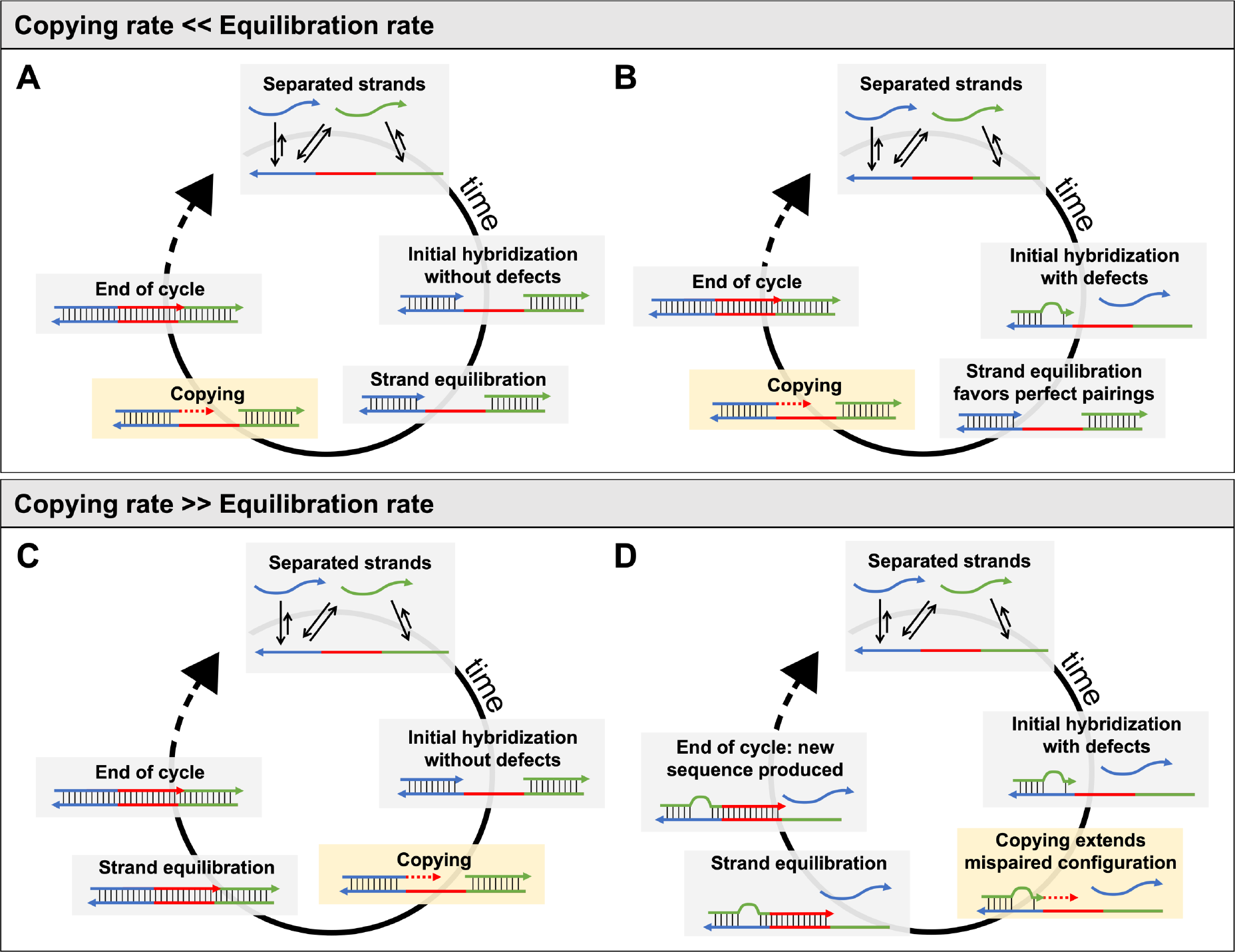
Copying of a template in a complex mixture. In a mixture of multiple RNA strands, short duplexes can dissociate because of chemical-physical changes such as dilution, thermal shock, reduction in ionic strength or change in pH (22, 60). As conditions favorable for pairing are re-established, complementary oligonucleotides quickly anneal. (**A, B**) If the equilibration rate is much faster than the copying rate, then regardless of the initial configuration of the duplexes, only the correctly paired primer will be elongated, and the information encoded in the template can be properly copied. (**C, D**) If the copying rate is faster than the equilibration rate, depending on the initial configuration it may be possible to either (**C**) properly copy the template or (**D**) to extend a mis-paired oligonucleotide, creating a novel sequence that was not previously encoded in the template.

Even a relatively small 11 nt long primer has a significant chance of finding an alternative binding site with two defects over a genome of 100nt in length (Supplementary Figure 8), a typical size for relevant ribozymes (61). We suggest that this physical mechanism may lead to loss of information even if the copying chemistry allowed for perfect fidelity (62). As we have shown (Figure 6A,B), this loss of fidelity due to incorrect primer/template association would be exacerbated by increasing the primer extension rate, suggesting the existence of an upper limit for a reliable copying rate, that is intimately linked to the rate of equilibration of the system (Figure 7).

In a biological context, given the generally accepted view that most of the human genome is transcribed (63), and given a total sequence length of ∼10^9^ nt, there is a significant chance (p > 0.05) that a random 20 nt oligonucleotide (a typical size for miRNA, ASO and siRNA) could hybridize to some cellular RNA with one or more mismatches or bulges. In a biological system, the transient low accuracy annealing that we have described may therefore exacerbate the extent of off-target hybridization by allowing the formation of undesired duplexes that would not be expected at equilibrium. These considerations suggest that over large genomes many imperfect duplexes that form by chance may persist in long-lasting out-of-equilibrium states, resulting in the dysregulation of gene expression and potential toxicity of oligonucleotide therapeutics (12, 64, 65). This in turn raises the question of how biological systems prevent or ameliorate such harmful non-specific interactions. We speculate that this may have contributed to the evolutionary selection of sequences with short-lived metastable states, or in the case of long RNA transcripts, the suppression of undesired associations through folding. Alternatively, the need to prevent deleterious mis-pairings may have led to the evolution of binding proteins that coat single stranded RNA and prevent mis-pairing from occurring (66), or RNA helicases that dissociate mis-paired RNAs.

Our analysis of the association and dissociation of imperfectly complementary oligonucleotides has led to surprising insights into the non-equilibrium properties of complex oligonucleotide mixtures. Our findings show the importance of appropriately accounting for mis-pairing interactions between oligonucleotides with mismatches and bulges when handling out-of-equilibrium mixtures of multiple strands. Future work will help to improve algorithms for the simulation of such mixtures (21), and will improve our understanding of RNA-RNA interactomes (67), facilitate the design of complex nucleic acid structures and help to predict the efficacy of gene and antisense therapies (68).

## ACKNOWLEDGEMENTS

We would like to thank Aleksandar Radakovic, Brennan Ashwood, Daniel Duzdevich and Filip Boskovic for comments on the manuscript.

## FUNDING

J.W.S. acknowledges support from the National Science Foundation [CHE-2104708 to J.W.S.], the Simons Foundation [290363]; the Sloan Foundation [19518], and the Moore Foundation [11479]. J.W.S. is an investigator of the Howard Hughes Medical Institute.

## CONFLICT OF INTEREST

None declared.

## Notes

### Competing Interest Statement

The authors have declared no competing interest.

### Summary of Updates

Main text and supplementary data have been revised.

