## Supplementary Data for "Transient states during the annealing of mismatched and bulged oligonucleotides"

### 1. List of sequences used

To perform the systematic characterization of the effect of defects on the annealing behavior of DNA and RNA, we designed a target (T) oligonucleotide 5′-TGGTGATGCGTG-3′ and edited its sequence introducing mismatches or bulges to produce competitors (C). All these sequences were tested for binding against a probe (P) of fixed sequence 5′-CACGCATCACCA-3. All target sequences and substitutions used for kinetic and thermodynamic characterization are listed below in Supplementary Table 1, with every sequence name referencing the nature and the position of the defects, either mismatches (m) or bulges (b), along the double helix.

| Name | Sequence | Dot-parens-plus (Probe+Sequence) | Defects listed from 5' to 3' | $k_{on}$ ($\mu M^{-1}s^{-1}$) | $K_D$ (M) |
|---|---|---|---|---|---|
| 7t | ATGCGTG | (((((((.....+))))))) | No defects | 9.71 ± 1.63 | $1.36 \times 10^{-6}$ |
| 8t | GATGCGTG | ((((((((....+)))))))) | No defects | 19.7 ± 1.01 | $1.28 \times 10^{-8}$ |
| 9t | TGATGCGTG | (((((((((...+))))))))) | No defects | 18.9 ± 2.86 | $2.59 \times 10^{-9}$ |
| 10t | GTGATGCGTG | ((((((((((..+)))))))))) | No defects | 18.3 ± 3.09 | $4.49 \times 10^{-11}$ |
| 11t | GGTGATGCGTG | (((((((((((.+))))))))))) | No defects | 21.5 ± 5.23 | $1.13 \times 10^{-12}$ |
| 12t | TGGTGATGCGTG | ((((((((((((+)))))))))))) | No defects | 26.6 ± 5.34 | $3.71 \times 10^{-13}$ |
| $m_{9,10}$ | TGAAGATGCGTG | ((((((((..((+))..)))))))) | G -> A, G -> A | 9.60 ± 2.76 | $5.19 \times 10^{-9}$ |
| $m_{6,7}$ | TGGTGTAGCGTG | (((((..(((((+)))))..))))) | A -> T, T -> A | 5.30 ± 0.17 | $4.46 \times 10^{-9}$ |
| $m_{4,9}$ | TGGAGATGAGTG | (((.((((.(((+))).)))).))) | T -> A, C -> A | 11.4 ± 0.59 | $2.39 \times 10^{-8}$ |
| $m_{10}$ | TGATGATGCGTG | (((((((((.((+)).))))))))) | G -> A | 15.0 ± 1.41 | $1.23 \times 10^{-10}$ |
| $m_7$ | TGGTGTTGCGTG | ((((((.(((((+))))).)))))) | A -> T | 16.4 ± 2.99 | $9.02 \times 10^{-11}$ |
| $m_{3,10}$ | TGATGATGCATG | ((.((((((.((+)).)))))).)) | G -> A, G -> A | 4.79 ± 0.76 | $8.27 \times 10^{-7}$ |
| $m_3$ | TGGTGATGCATG | ((.(((((((((+))))))))).)) | G -> A | 10.4 ± 1.20 | $1.44 \times 10^{-9}$ |
| $m_{3,4}$ | TGGTGATGTATG | ((..((((((((+))))))))..)) | C -> T, G-> A | 18.0 ± 2.75 | $2.23 \times 10^{-8}$ |
| $m_5$ | TGGTGATACGTG | ((((.(((((((+))))))).)))) | G -> A | 14.4 ± 2.72 | $9.89 \times 10^{-10}$ |
| $m_{12}$ | AGGTGATGCGTG | (((((((((((.+.))))))))))) | T -> A | 21.9 ± 2.42 | $2.04 \times 10^{-12}$ |
| $m_1$ | TGGTGATGCGTC | .(((((((((((+))))))))))). | G -> C | 11.2 ± 0.81 | $6.67 \times 10^{-13}$ |
| $m_{1,12}$ | AGGTGATGCGTC | .((((((((((.+.)))))))))). | T -> A, G -> C | 12.0 ± 1.44 | $9.15 \times 10^{-13}$ |
| $b_{10}$ | TGTGATGCGTG | (((((((((.((+))))))))))) | G -> removed | 14.8 ± 1.15 | $3.39 \times 10^{-11}$ |
| $b_8$ | TGGTATGCGTG | (((((((.((((+))))))))))) | G -> removed | 14.7 ± 1.08 | $6.73 \times 10^{-9}$ |
| $b_6$ | TGGTGAGCGTG | (((((.((((((+))))))))))) | T -> removed | 22.2 ± 3.42 | $2.18 \times 10^{-10}$ |
| $b_4$ | TGGTGATGGTG | (((.((((((((+))))))))))) | C -> removed | 29.7 ± 5.27 | $9.17 \times 10^{-9}$ |
| $b_2$ | TGGTGATGCGG | (.((((((((((+))))))))))) | T -> removed | 17.0 ± 2.17 | $7.97 \times 10^{-12}$ |
| $12t^{RNA}$ | UGGUGAUGCGUG | ((((((((((((+)))))))))))) | No defects | 32.5 ± 4.46 | $2.89 \times 10^{-19}$ |
| $m_7^{RNA}$ | UGGUGUUGCGUG | ((((((.(((((+))))).)))))) | A -> U | 32.0 ± 1.44 | $8.82 \times 10^{-15}$ |
| $m_{4,9}^{RNA}$ | UGGAGAUGAGUG | (((.((((.(((+))).)))).))) | U -> A, C -> A | 12.0 ± 1.44 | $1.66 \times 10^{-9}$ |
| $m_{6,7}^{RNA}$ | UGGUGUAGCGUG | (((((..(((((+)))))..))))) | A -> U, U -> A | 3.65 ± 3.70 | $2.87 \times 10^{-11}$ |
| $b_6^{RNA}$ | UGGUGAGCGUG | (((((.((((((+))))))))))) | U -> removed | 29.8 ± 4.09 | $1.56 \times 10^{-12}$ |

**Supplementary Table 1.** List of sequences used in this work and their relative measured parameters. The subscript in the sequence name refers to the position of the defects along the probe sequence as defined in the main text. Data for control RNA sequence ($12t^{RNA}$) is from our recent work (1).

#### 2. Measurement of hybridization rates and dissociation constants

To determine the hybridization rates for the oligonucleotides studied in this work, the fluorescent traces for 2-aminopurine emission have been collected performing stopped-flow experiments and individually fitted to find the optimal $k_{on}$. For our fitting routine, simulated traces have been calculated by numerically integrating the following set of coupled differential equations using the variable step solver *ODE15s* in MATLAB:

$$\begin{cases} \frac{d[A]}{dt} = -[A]k_{on} + [AB]k_{on}K_D \\ \frac{d[B]}{dt} = -[A]k_{on} + [AB]k_{on}K_D \\ \frac{d[AB]}{dt} = +[A]k_{on} - [AB]k_{on}K_D \end{cases}$$

(Supplementary Eq. 1)

The mean squared residuals with respect to our experimental data have been minimized using MATLAB *fminsearch* function to determine the best fitting $k_{on}$, yielding the overlapping traces shown in the example in Supplementary Figure 1A. The $k_{on}$ values computed this way have been averaged to obtain the final values tabulated in Supplementary Table 2. $K_D$ values used to calculate hybridization traces and $k_{off}$ have been determined through a series of melting experiment as described in the main text, yielding the fluorescent traces shown in the example in Supplementary Figure 1B. These traces have been individually fitted to extract estimates and confidence intervals of $K_D$ at 25°C.

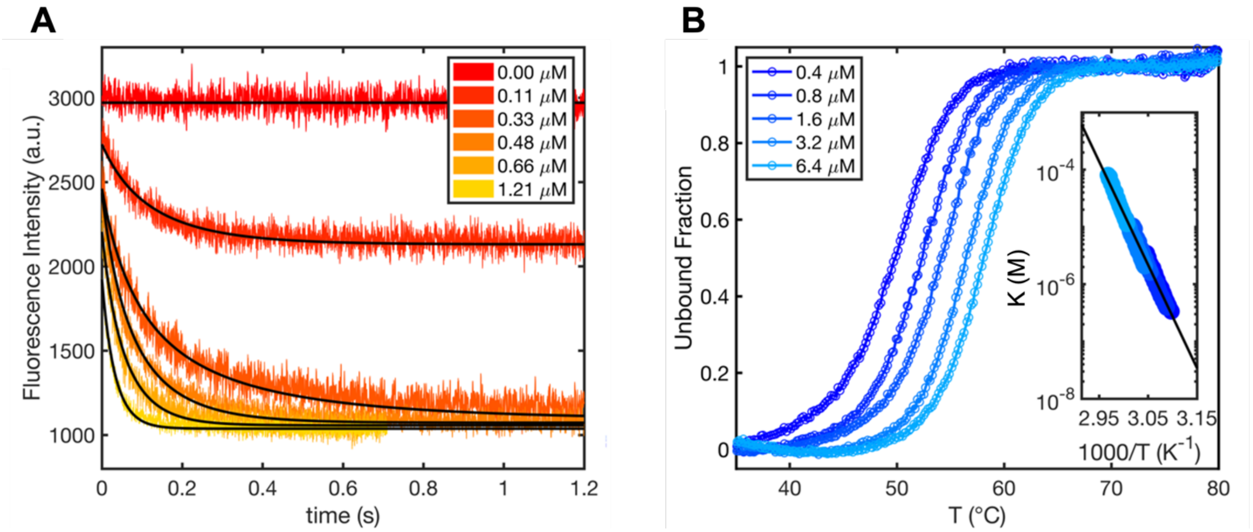

**Supplementary Figure 1.** (**A**) Examples of raw stopped-flow data traces and best-fit simulation traces. (**B**) Examples of fluorescence melting curves and the corresponding van't Hoff plot in the inlay.

#### 3. Fitting analytical model for hybridization

In order to separate the contributions of defects into (i) loss of nucleation sites and (ii) interference in the zippering process, we implemented an analytical model recently presented by Hertel and colleagues (2). The model is controlled by two free parameters, namely the bimolecular nucleation rate and the ratio ($\alpha$) between the zippering rate of the duplex (following the formation of the nucleus) and the dissociation rate of the nucleation region.

Based on the definition of $p^{zippering}$, we can intuitively see how this value is maximally sensitive to $\alpha$ in a range dependent on the free energy change upon the formation of a nucleation region.

We can conveniently calculate the theoretical lower and higher boundaries for $\alpha$ that we are sensitive to, depending on the nucleation sites in our database and user-defined probability thresholds. These thresholds effectively bound the $p^{zippering}$ function between $th_{low}$ and $th_{high}$, that we have arbitrarily set as equal to 0.01 and 0.99, meaning that we are discarding the infinite values of $\alpha$ that would either decrease $p^{zippering}$ below 0.01 or increase it above 0.99 and that cannot reasonably be distinguished experimentally. These experimental boundaries can be determined as follows:

$$\begin{cases} \alpha_{min} = \dfrac{th_{low} \cdot e^{\Delta G^{min}/RT}}{1 - th_{low}} = \dfrac{0.99 \cdot e^{-16.1}}{0.01} \approx 10^{-9} \\ \alpha_{max} = \dfrac{th_{high} \cdot e^{\Delta G^{max}/RT}}{1 - th_{high}} = \dfrac{0.01 \cdot e^{-8.8}}{0.99} \approx 1.5\ x\ 10^{-2} \end{cases}$$

(Supplementary Eq. 2)

In our fitting routine, the squared sum of the residuals was minimized using MATLAB *fminsearchbnd* function (3) in order to obtain a single shared $k_{nuc}$ and 4 different bounded $\alpha$ values for either perfectly paired duplexes, or duplexes with mismatches, double mismatches or bulges. To improve the robustness of the minimization process, the logarithms of all 5 free parameters were passed to *fminsearchbnd*.

Finally, we used a Monte Carlo method to estimate confidence intervals for our parameters: we resampled the dataset 1000 times, drawing each $k_{on}$ value from a normal distribution defined according to its experimentally determined error range.

For each iteration of the fitting routine, we checked the curvature of the objective function at its minimum to evaluate how well constrained the $\alpha$ values were. While for mismatched and double mismatched duplexes these values were well constrained, for perfect and bulged duplexes we could reliable estimate only a lower boundary (set at 99% of the minimum of the objective function). These values are reported in Supplementary Figure 2 (panels A-D). For each iteration, we could use the fitted $\alpha$ value to compute an associated distribution of $p^{zippering}$, whose aggregated values distributions are shown in Supplementary Figure 2 (panels E-H).

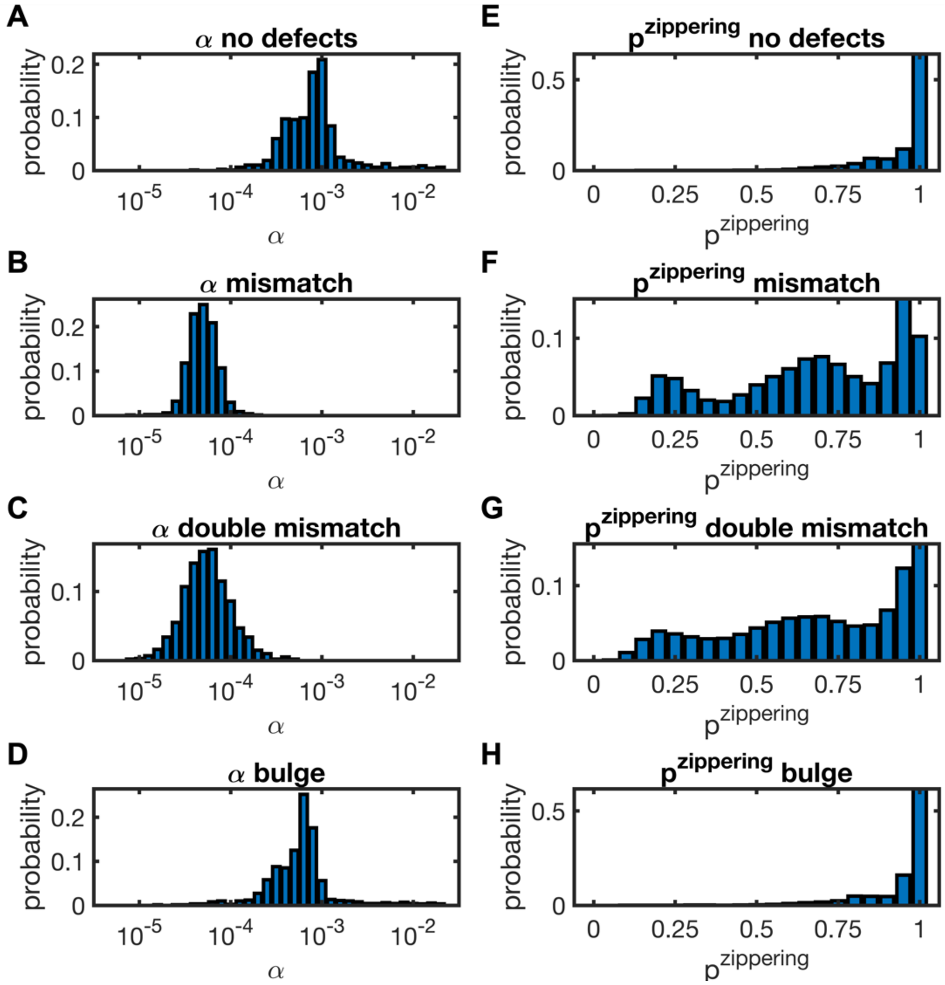

**Supplementary Figure 2.** (**A-D**) Best fitting *α* values determined in our Monte Carlo sampling procedure. (**E-H**) Probability of successfully zippering upon nucleus formation determined using the aggregated *α* values.

Median values and associated confidence intervals for the parameters obtained in this work are tabulated below in Supplementary Table 2:

| Parameter | Median value | CI (95%) |
|---|---|---|
| $k_{nuc}$ | $2.2 \times 10^{6}$ $M^{-1}s^{-1}$ | $(1.9 – 2.6) \times 10^{6}$ $M^{-1}s^{-1}$ |
| $\alpha^{perfect}$ | $7.2 \times 10^{-4}$ * | $1.3 \times 10^{-4} – 6.5 \times 10^{-3}$ * |
| $\alpha^{mismatch}$ | $4.3 \times 10^{-5}$ | $2.2 \times 10^{-5} – 8.9 \times 10^{-5}$ |
| $\alpha^{double\ mismatch}$ | $4.6 \times 10^{-5}$ | $1.6 \times 10^{-5} – 1.7 \times 10^{-4}$ |
| $\alpha^{bulge}$ | $5.4 \times 10^{-4}$ * | $8.3 \times 10^{-5} – 5.6 \times 10^{-3}$ * |

**Supplementary Table 2.** Coefficients determined from global fitting of the analytical model.
***** *α* for perfect duplexes and duplexes with bulges should be treated as lower boundaries.

### 4. Effect of defects on RNA kinetics and generalization

We have established that almost all destabilization coming from mismatches and bulges in a DNA duplex comes from changes in dissociation kinetics, with association rates being affected to a minor extent. We have addressed the same issue in RNA duplexes by studying the behavior of a subset of the sequences used in our DNA studies. Our RNA results are shown in Supplementary Figure 3 and show the same behavior as observed for DNA. We find a comparable penalty for each studied defect in DNA and RNA.
From this study we see that the ΔG of hybridization is well correlated with association rates for perfectly paired duplexes ($R^2$ = 0.76), while this is not the case for defect-bearing duplexes ($R^2$ = 0.08) as shown in Supplementary Figure 3C. This difference highlights the fact that defects affect annealing process in a non-trivial way. The opposite is true for the dissociation of the two strands, so that the logarithm of the dissociation rate is very well correlated with ΔG of hybridization both for perfectly pairing and defect-bearing duplexes as shown in Supplementary Figure 3D.

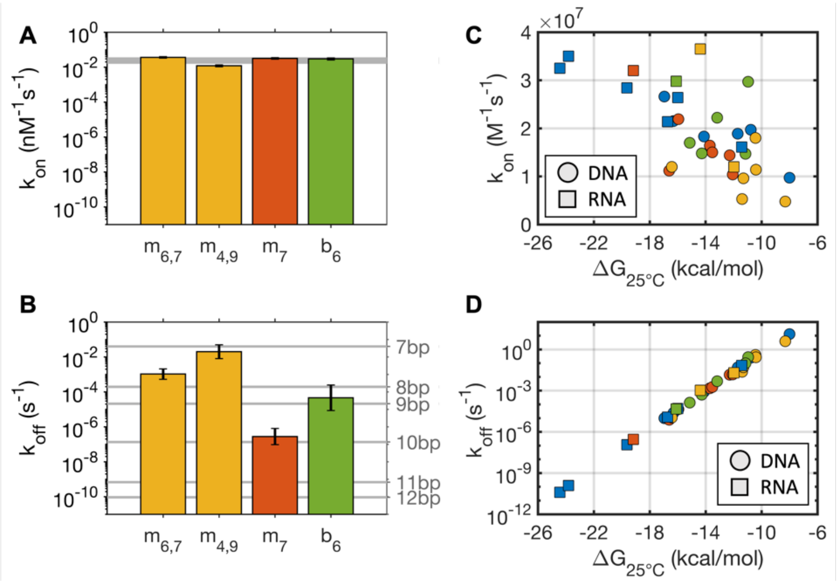

**Supplementary Figure 3.** Association and dissociation kinetics for short RNA duplexes with mismatches and bulges. (**A**) Association and dissociation rate constants for RNA oligonucleotides with either single mismatches, bulges or double mismatches. The grey shaded band highlights typical $k_{on}$ values for control oligonucleotides with length varying from 7bp to 12bp, yielding values varying by roughly a factor of two. Data are shown on a logarithmic scale to directly compare the relative contributions of $k_{on}$ and $k_{off}$ to $K_D$. (**B**) Dissociation rate constants for short RNA oligonucleotides with either single mismatches, double mismatches, or bulges. Horizontal lines refer to $k_{off}$ for a series of perfectly pairing control duplexes. All data for perfectly pairing RNA duplexes have been taken from the literature (1). (**C, D**) Overview of $k_{on}$ (**C**) and $k_{off}$ (**D**) as a function of hybridization energy for DNA and RNA with and without defects.

To further extend and generalize our study on RNA, we measured hybridization rates for a biologically relevant sequence (miR125, rUrCrCrCrUrGrArG) in a buffer that more closely resembles physiological conditions.
Micro RNAs (miRNA) are short (20-24 nt) oligonucleotides known to regulate eukaryotic gene expression. Animal miRNAs act by pairing their 5' "seed" region to the 3' UTR region of target messenger RNA transcripts and inducing translational inhibition, accelerated exonucleolysis and slicing (4–6). *In vivo* studies have shown that the pairing of nucleotides 2-7 of the miRNA is often necessary to correctly induce mRNA regulation, leading to the formulation of the so-called "seed rule". This rule – not without exceptions – is so powerful as to be used as a guide for tools predicting putative miRNA target sites (6–8). While the mechanism that lies behind this empirical rule is not clear, following the approach by Cisse et al. (9) we tested the effect of mismatches in the 8bp long "seed" sequence of the human miR125. We mutated the target sequence in position 2, reducing the length of the longest continuous stretch down to 6bp. The effect of this mutation is that such sequence would not be recognized as a target anymore according to the "seed

rule”. One possible explanation for such phenomenon, would be that reducing the length of the “seed”, some intrinsic physical feature of the hybridization could be affected, a hypothesis that we tested by performing a series of hybridization experiments in physiological salt conditions (200mM NaCl, 10mM Tris pH 8.0) as shown in Supplementary Figure 4. Our results show no significant difference in the association rate between the wild-type and seed-less duplex.

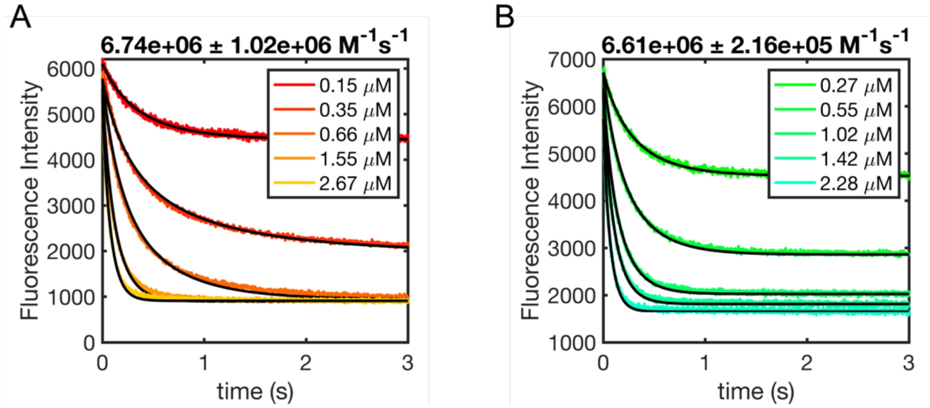

**Supplementary Figure 4.** (**A**) Time traces for the hybridization of the human miR125 seed sequence to its perfectly complementary sequence (rCrUrCrArGrGrGrA) and (**B**) its variant (rCrUrCrArGrGrUrA) bearing a single mismatch in 200mM NaCl, 10mM Tris pH 8. The miR125 seed sequence containing 2Ap was used at a fixed concentration of 0.43 μM, while the perfectly complementary sequence and mismatched variant sequence concentrations were varied according as specified in the inserts in panel A and B respectively. Values above panels reflect the determined association rates and relative errors for the two case studies.

#### 5. Comparison with NN predictions

In our study we have found that oligonucleotide dissociation rates are well-correlated with free energy change upon hybridization. A good agreement between the latter and NN predictions would make them a powerful tool to quickly calculate dissociation rates for any arbitrary duplex sequence with defects. We show here the direct comparison for measured and predicted ΔG at 25°C, with results shown in Supplementary Figure 5.

We find an excellent correlation between the two values ($R^2$ = 0.95) with a discrepancy of +0.9 kcal, which is readily explained by the duplex destabilization due to the 2-aminopurine used in our experiments.

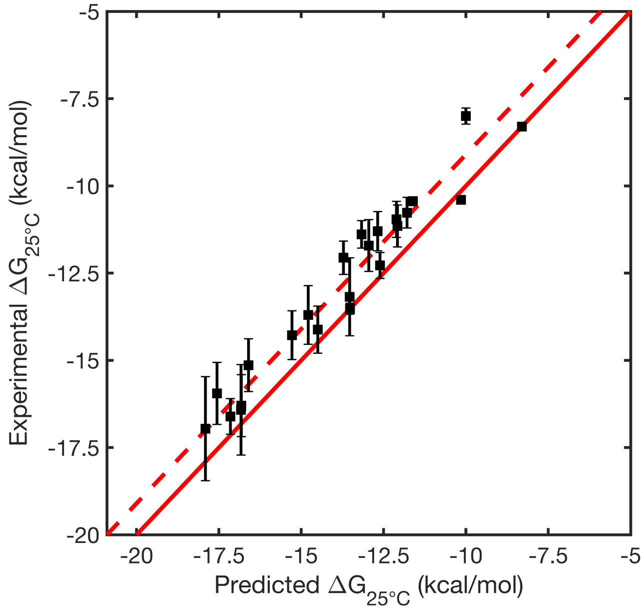

**Supplementary Figure 5.** Comparison between predicted and measured ΔG at 25°C for the DNA oligonucleotides used in this study. Continuous line shows perfect agreement between the two datasets, while dashed line corresponds to predictions when the penalty from 2-aminopurine is taken into account.

#### 6. Equilibration of competing RNA mixtures

Since we have found comparable association rates for perfectly pairing, mismatched and bulged RNA duplexes, we moved on to study the equilibration of mixtures of competing oligonucleotides. We found that DNA mixtures show a transient low-accuracy state upon mixing, and we decided to test whether the same was true for RNA. We performed a series of competition experiments between equimolar amounts of probe sequence (P), perfectly pairing target (T) and defected competitor (C), with target strands bearing a fluorescent nucleotide (2Ap) to report the bound fraction of T over time. The results of this study are shown here in Supplementary Figure 6, where the fluorescent signal has been normalized to yield 1-$\theta$, with $\theta$ defined in the main text as equal to *[T:P] / ([T:P] + [C:P]).*

When working with RNA, since the target-competitor mixtures contain high GU-content sequences, before mixing with the probe these were briefly heated up to 90°C for 3 minutes to remove any potential structures inherited from their highly concentrated stock solutions.

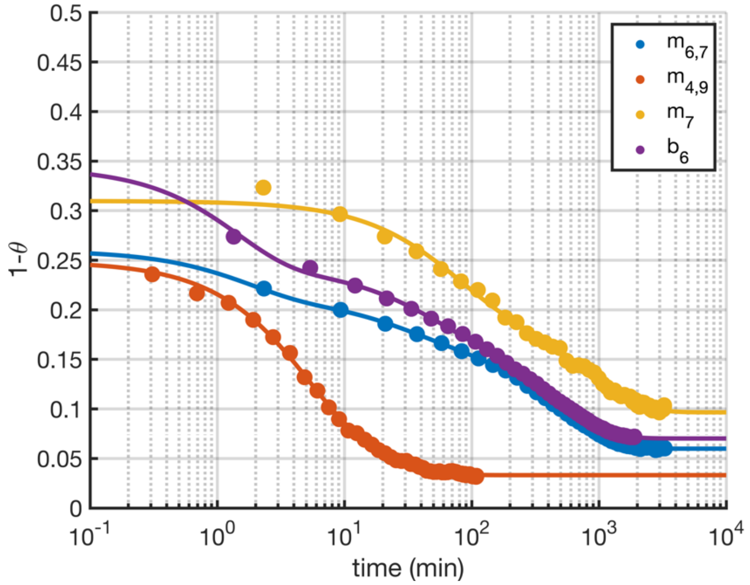

**Supplementary Figure 6.** Equilibration in competing mixtures of short RNA oligonucleotides. In each experiment shown here, equimolar amounts of probe sequence (P), perfectly pairing target (T) and mismatched or bulged competitor (C) were mixed at a final concentration of 1 µM, and the fluorescent emission of T was tracked. The competitor used in each mixture is noted in the inset.

### 7. Kinetics of non-enzymatic primer extension and ligation in competing mixtures

To evaluate whether defect-containing duplexes that are virtually absent at equilibrium could be substrates for relevant non-enzymatic chemical reactions, we performed a set of primer extension and ligation experiments in a competitive scenario using a perfectly pairing RNA target and a double mismatched RNA competitor.
To make the target and competitor strands suitable substrates for these chemical reactions, the sequence design of the probe had to be modified to present an overhang. All sequences used in these experiments are listed in Supplementary Table 3.

| Name | Sequence (5′→3′) |
|---|---|
| Probe for primer extension | GGGG CAC GCA UCA CCA |
| Complement to the probe for primer extension | G UGG UGA UGC GUG CCCC G |
| Probe for ligation | UGCG CAC GCA UCA CCA |
| Complement to the probe for ligation | G TGG TGA TGC GTG CGCA G |
| Perfectly matched target ($12t^{RNA}$) | 5′-FAM-UGG UGA UGC GUG |
| Mismatched competitor ($m_{4,9}^{RNA}$) | 5′-CY5-UGG AGA UGA GUG |

**Supplementary Table 3.** List of RNA sequences used for non-enzymatic primer extension and ligation.

To monitor the extent of chemical primer extension and ligation, the competing oligonucleotide C was labelled with a 5′-Cy5 fluorescent dye, while the target oligonucleotide T was labelled with a 5′-FAM fluorescent dye. Denaturing polyacrylamide gels were imaged in the green and red channels to quantify the reacted target strand and the reacted competitor strand over time, as depicted in the example in Supplementary Figure 7.

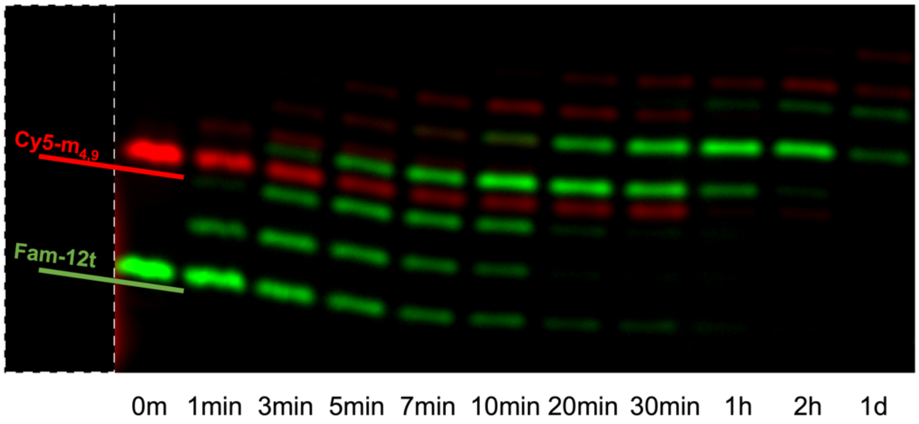

**Supplementary Figure 7.** Image of a denaturing PAGE used to resolve reaction products in a competition experiment in presence of 1.5 mM C*C. The red and green channels have been overlaid to show the reaction products of both species as a function of time. Contrast has been enhanced for visualization purposes.

#### 8. Calculation of mismatch probability

To quantify the extent at which mismatched sequences could affect prebiotic or biological processes in an out-of-equilibrium scenario, we needed first to determine how probable it is to have mismatch or bulged pairings by chance between two strands. The probability $P_{pair}$ of a random $N$-nt long sequence matching a given target with a number $d$ of defects (mismatches, bulges…) is equal to:

$$P_{pair}(N,d) = \left(1-\frac{1}{4}\right)^{d} \cdot \left(\frac{1}{4}\right)^{N-d} \cdot \binom{N}{d}$$

(Supplementary Eq. 3)

where $\binom{N}{d}$ is the binomial coefficient that represents the number of ways to arrange the defects across the $N$ bases of the duplex. The predictions from this simple function are shown in Supplementary Figure 8A, showing that the probability of pairing for two random oligonucleotides increases as more defects are introduced until a maximum value is reached, depending on the total length of the strands.

This function only applies to the case when two and only two sequences are present in the same environment, which is not the case we were interested in. Instead, we wanted to know how likely it is for a given sequence to find a target pairing partner in an environment containing a large set of oligonucleotides or long strands with a multitude of subsequences available for pairing. Here we refer to this set of potential pairing partners as a genome, but this ‘genome’bcould be either be physical oligonucleotides of length $L$ (such as a biological genome) or a collection of oligonucleotides whose total length is equal to $L$ (such as a potentially prebiotic genome). The probability $P_{genome}$ of observing at least one occurrence of a potential pairing partner for a strand of length $N$ with $d$ mismatches or bulges in a genome of length $L$ is then equal to:

$$P_{genome}(L,N,d) = 1 - \left[1 - P_{pair}(N,d)\right]^{K(L)}$$

(Supplementary Eq. 4)

with $K(L)$ equal to $2L$ for a circular genome or $2(L\text{-}N\text{+}1)$ for a linear genome, when both sense and antisense strands are considered. Using these functions, it is possible to compute the probability of off target binding in a genome as defined above (Supplementary Figure 8B). Given a random oligonucleotide of length $N$, the longer the genome and the higher the number of defects, the higher the probability of finding a pairing site by chance.

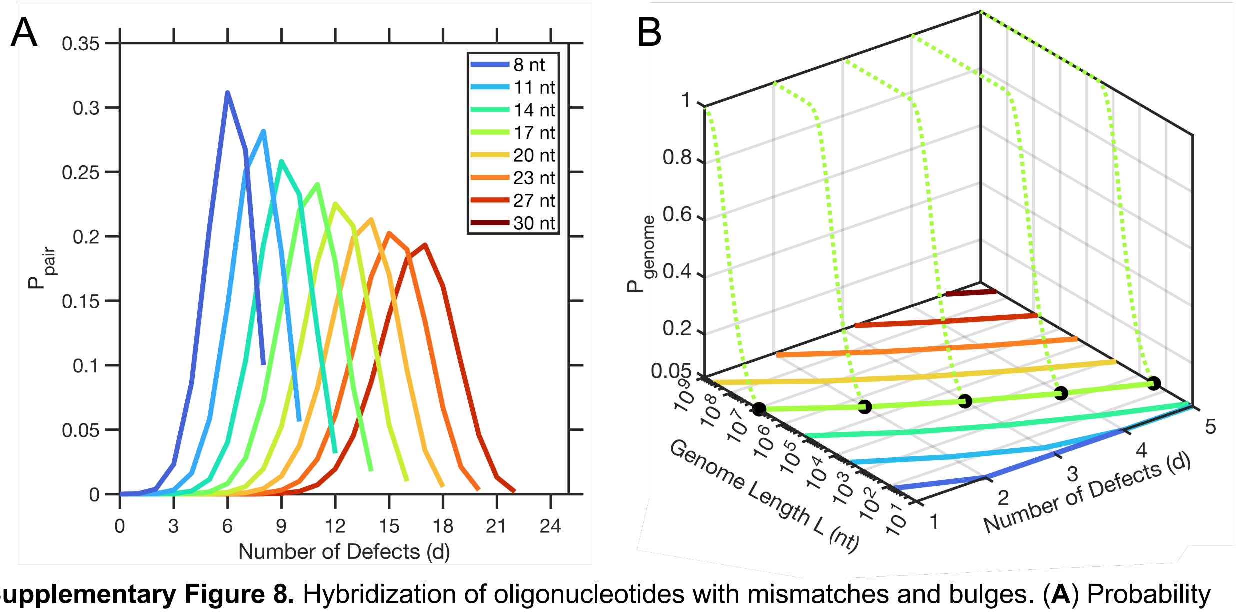

**Supplementary Figure 8.** Hybridization of oligonucleotides with mismatches and bulges. (**A**) Probability of two random strands forming a duplex with defects. The function is not monotonic, since partial complementarity between two strands is more likely than both perfect complementary or total non-complementary. (**B**) Relationship between the length of a genome $L$ and the probability $P_{genome}$ of finding a potential pairing site with d defects for a 17nt long oligonucleotide (dotted green lines). If we arbitrarily pick a threshold probability of 0.05 for such a pairing site, we can trace continuous lines on the 2D plane representing the minimum genome length at which the binding of an oligonucleotide with *d* defects becomes relevant ($P_{genome}$ > 0.05)
